# Preserving Nanotherapy-Induced Allogeneic Murine Islet Graft Tolerance During Distal Methicillin-Resistant Staphylococcus aureus Skin Infection

**DOI:** 10.64898/2026.11.04.763192

**Authors:** El Hadji Arona Mbaye, Débora B. Scariot, Xiaomin Zhang, Calleigh P. Reber, Natalie R. Klug, Szumo Wang, Sultan Almunif, Amanda B. K. Bakoshi, Rayanne R. M. Machado, Natalia Mendoca, Andrew J. Monteith, Evan A. Scott, Jacqueline A. Burke

## Abstract

Eliminating lifelong immunosuppression can make islet transplantation a durable alternative to insulin therapy for type 1 diabetes. As short-course tolerogenic regimens are developed to replace lifelong immunosuppression, it is critical to test their robustness against perturbations such as hospital-acquired infections, which transplant recipients routinely encounter. Herein, we investigated whether rapamycin-loaded polymersomes (rPS), a subcutaneous nanotherapy that induces long-term murine islet allograft survival, can preserve host defenses in a Methicillin-resistant *Staphylococcus aureus* (MRSA) skin infection, and whether the resulting immune response from this distal infection relative to graft location erodes rPS-induced tolerance. rPS maintained neutrophil antibacterial activity despite inhibiting oxidative mechanisms *in vitro*, and rPS-treated mice demonstrated normal recovery following MRSA challenge. However, MRSA skin infection caused a decline of ∼43% in recipient survival for rPS-treated mice, despite surviving recipients retaining glucose-stimulated insulin responsiveness. This erosion of allograft acceptance coincided with a burst of IL-6 within 24h of the infection followed by elevated TNF-□ and neutrophilia at 13 days post-infection. Combining rPS with a single dose of methylprednisolone reduced IL-6 and TNF-α without altering neutrophilia, nearly doubling survival among infected recipients to 80%. Our findings demonstrate that partial dampening of inflammatory insults can restore superior islet allograft acceptance following perioperative infection.

## Introduction

Transplantation (TX) remains dependent on perpetual, non-specific immunosuppression that is often toxic and associated with severe side effects.^1^ In a recently published survey by the American Society of Transplantation, 92% of respondents have reported side effects attributable to lifelong immunosuppression which increases susceptibility to infections leading to increased graft failure and mortality.^2,3^ As such, the field has long aimed to move away from lifelong immunosuppression and instead achieve antigen-specific tolerance (AST). AST is defined as unresponsiveness to a particular antigen in the graft long after discontinuation of a short-treatment course without global immunosuppression. Tolerance induction has converged on a limited set of cellular mechanisms including hyporeactivity or anergy of alloreactive T cells and pronounced regulatory T cells (Tregs) expansion, both of which can be induced through costimulation blockade via the use of monoclonal antibodies or targeted delivery of small molecules to antigen-presenting cells (APCs).^4–9^ While AST remains clinically unachieved, it has been realized in murine models as we have previously demonstrated using subcutaneously administered rapamycin-loaded polymersomes (rPS) nanotherapy. We found that rPS achieves 100-day normoglycemia in 83% diabetic mice receiving allogeneic islet transplants by inducing downregulation of costimulatory markers, CD4+ T cell anergy, and CD8+ Treg expansion in the axial and brachial lymph nodes.^9^ Furthermore, rPS-treated recipient T cells were unresponsive to donor antigen while remaining responsive to third-party antigen, establishing AST establishment in this model.^9^

While these findings substantiate the translational potential of rPS-induced AST, achieving durable tolerance is only half of the challenge; the other lies in making it resilient. With the goal of realizing “one transplant for life”, the clinical robustness of AST-inducing therapies will depend on their ability to withstand nosocomial infections, a perturbation that transplant recipients routinely encounter during the same window in which tolerance is being established.^10,11^ Skin and soft tissue infections account for roughly 20% of nosocomial infections, and Methicillin-resistant *Staphylococcus aureus* (MRSA) in particular is among the most common pathogens encountered in the first month after TX, precisely when tolerance is most vulnerable to disruption.^11,12^ TX tolerance is increasingly recognized not as a binary, all-or-none state but as a graded phenomenon whose robustness depends on the number and redundancy of underlying cellular mechanisms engaged during its induction.^13^ Because tolerance can rest on multiple, partially overlapping pathways, a state that appears fully tolerant by clinical or histological criteria may still vary in its underlying resilience, making it differentially susceptible to perturbations.^13^ Bacterial infections have been shown to disrupt this metastability, overriding costimulation-blockade-induced tolerance through distinct cytokine pathways that reactivate alloreactive T cells.^14–17^ Previous work on erosion of TX tolerance due to infection has mainly focused on systemic infections, raising the question whether a distal skin infection relative to graft location can generate similar magnitudes of tolerance disruption.^14,16,18^

To evaluate this vulnerability, we superimposed a murine MRSA skin infection on rPS-induced islet allograft tolerance. We hypothesized that redirecting rapamycin toward antigen-presenting cells would spare neutrophil-mediated antibacterial defense, and that a cutaneous infection anatomically distant from the islet graft would spare allograft acceptance. Herein, rPS preserved neutrophil bactericidal capacity and viability despite suppressing oxidative antibacterial mechanisms, yet the distal lesion site infection still eroded long-term allograft acceptance. Erosion tracked with a transient serum IL-6 increase and lymphopenia within 24 hours of the infection later followed by increased TNF-□ and elevated neutrophil counts. Blunting pro-inflammatory cytokines with a single dose of methylprednisolone restored graft acceptance without eliminating early lymphodepletion or neutrophilia in the blood. As such, infection-driven tolerance loss emerged as a graded, time-dependent process in which reducing an early inflammatory cascade, rather than total inflammatory burden, can sustain the tolerant state after subcutaneous MRSA infection.

## Materials & Methods

### 2.1. MRSA infection study

Healthy C57BL/6 mice (9-12 weeks old) were dosed with subcutaneous rPS every 3 days for 6 doses (rPS 6x + MRSA) starting at –7 days post-infection (dpi). Experimental controls included infected (MRSA), uninfected (Control) groups and mice receiving Rapamune oral solution (VistaPharm) daily for 14 days (Rapamune PO 14x + MRSA) as a control of standard systemic immunosuppression. All drug formulations were diluted to 0.125 mg rapamycin /mL in PBS and administered at a dosage of 1 mg rapamycin/kg. On day 0, infected groups received a subcutaneous inoculation of MRSA strain JE2 (10^9^ CFUs in 100 μL) in the dorsal region. Body weight and lesion area were monitored throughout the study. Lesion areas were quantified in ImageJ using the average lesion length and average width.^25^ Serum was collected at 3 dpi and at endpoint (10 dpi) for cytokine analysis. Upon euthanasia, liver, and skin were harvested for bacterial burden analysis, and the skin was additionally processed for cytokine and myeloperoxidase (MPO) levels. Spleens were collected for flow cytometry.

### 2.2. Allogeneic islet TX + MRSA infection study

Five days prior to islet TX, diabetes was induced in C57BL/6 recipients by streptozotocin (STZ) administration (190 mg/kg) and confirmed by two consecutive nonfasting blood glucose (BG) >300 mg/dL. At –2 dpi, mice received Rapamune by oral gavage or subcutaneous rPS every 3 days for 6 doses (1 mg rapamycin/kg). BALB/c islets were isolated via collagenase digestion and density purification and transplanted under the kidney capsule of diabetic recipients. One day after TX, infected groups received a subcutaneous dorsal inoculation of MRSA strain JE2 (5 × 10^8^ CFUs in 100 μL). Weight, BG and lesion area were monitored for up to 59 dpi. Blood was collected at –8 dpi and post-infection (16 hours post-infection (hpi), 1, 13, and 29 dpi) for cytokine analysis and complete blood count. An intraperitoneal glucose tolerance test (IPGTT) was performed on surviving recipients at 27 and 57 dpi. Mice were fasted for 12h prior to testing, then administered dextrose (2g/kg) intraperitoneally, and BG levels were monitored at 0, 10, 20, 30, 60, 90, and 120 mins post-injection.

### 2.3. Additional Methods

Please consult Supplementary Methods.

## Results

### 3.1 Characterization of rPS

rPS were formulated with an average diameter 110.0 ± 1.0 nm and a polydispersity index (PDI) of 0.123 ± 0.016, demonstrating a monodisperse formulation (Fig. 1A, D). rPS matched the bilayer vesicle structure obtained from small-angle X-ray scattering which was visually confirmed by cryogenic transmission electron microscopy (Fig. 1B, C). Rapamycin encapsulation efficiency within polymersomes was 58.6%, and endotoxin levels were 0.012 EU/mL (Fig. 1D), below the immune response criterion.^26^ Blank PS possessed a size of 113.4 ± 1.7 nm and a PDI of 0.133 ± 0.043 (Fig. 1D). Hence, rPS and blank PS showed no significant differences in size and morphology.

**Figure 1.**
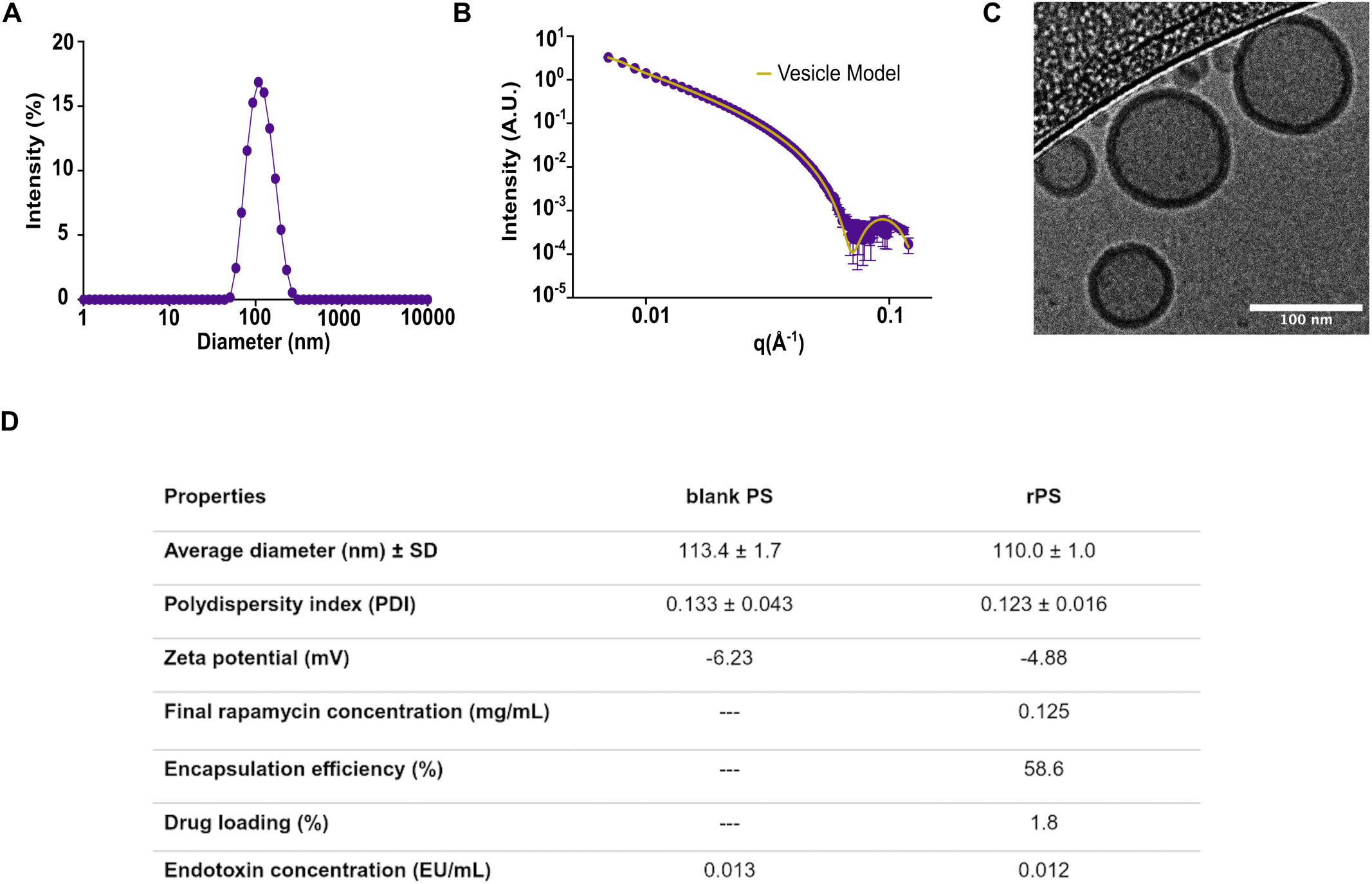
Characterization of engineered rapamycin-loaded polymersomes (rPS) and blank polymersomes (blank PS). (A) Size distribution of rPS as measured by DLS. (B) SAXS analysis of rPS fitted to a polymeric vesicle model using SasView software. (C) Cryo-TEM image verifies the bilayer vesicular structure of rPS. (D) Summary of the characterization parameters of rPS and blank PS, including size (nm), polydispersity index (PDI), zeta potential (mV), final rapamycin concentration (mg/mL) of rPS for *in vivo* experiments, encapsulation efficiency (%), drug percent loading (%) and endotoxin concentration (EU/mL).

### 3.2 rPS-treated neutrophils maintain antibacterial activity

Neutrophils are prominent players in the response against MRSA given their ability to trigger potent oxidative and non-oxidative killing mechanisms to guarantee pathogen control.^27^ Hence, to understand the effect of rPS on neutrophils’ bacterial killing, neutrophils were isolated from the femurs and tibia of healthy mice and treated with rPS, rapamycin, or blank PS. rPS was similar to rapamycin and inhibited reactive oxygen species production in neutrophils following lipopolysaccharide stimulation compared to the stimulated control (Fig. 2A). Blank PS produced an equivalent reduction in ROS (p < 0.001 vs Stim) highlighting that the poly(propylene sulfide) block of the poly(ethylene glycol)_17_-b-poly(propylene sulfide)_30_ (PEG_17_-*b*-PPS_30_) polymer, a known ROS scavenger,^28^ is itself sufficient to downregulate oxidative mechanisms. Given the reliance of NETosis on oxidative processes,^29^ suicidal NETosis was equally inhibited by rPS, rapamycin and blank PS (Fig. 2B). However, only rapamycin significantly inhibited the antibacterial activity of neutrophils whereas rPS showed 25% bacterial survival respectively similar to the control (Stim) (Fig. 2C). This dissociation between bacterial survival and oxidative mechanisms was not attributable to neutrophil toxicity from rPS or blank PS, as neutrophil viability was preserved in both groups. In contrast to rPS, rapamycin triggered significant neutrophil death (p < 0.0001) (Fig. 2D). Taken together, these results show that rPS inhibits oxidative mechanisms of neutrophils while preserving both neutrophil viability and non-oxidative antibacterial killing, a pattern not observed with rapamycin.

**Figure 2.**
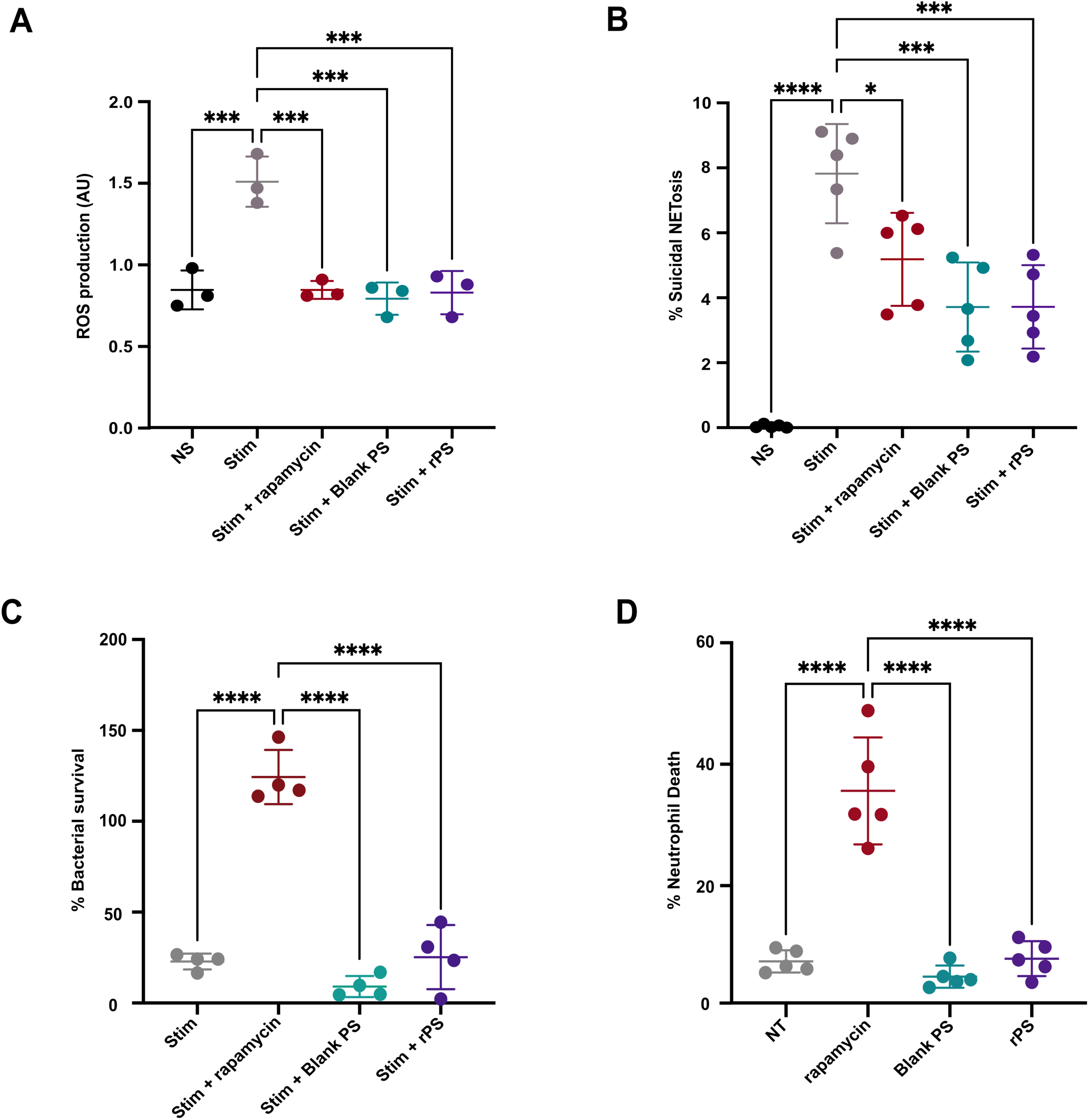
rPS preserves antibacterial activity of murine neutrophils *in vitro*. (A) Total reactive oxygen species (ROS) produced by lipopolysaccharide (LPS)-stimulated neutrophils isolated from the tibias and femurs of mice following treatment with rapamycin, blank PS, or rPS. Untreated and unstimulated cells were labelled “NS” and untreated stimulated cells “Stim” served as controls stimulated with LPS. (N = 3/treatment). (B) Percent (%) suicidal NETosis in *S.aureus* JE2 strain stimulated mouse neutrophils following the same treatment groups as in (A). Suicidal NETosis was measured via flow cytometry with staining of extracellular dsDNA, MPO and H3Cit. (N = 5/treatment). (C) Survival of *S. aureus* JE2 strain culture with treated mouse neutrophils. (N = 4/treatment). (D) Percent neutrophil death in mouse neutrophils treated with rapamycin, blank PS and rPS measured via flow cytometry by live/dead staining. “NT” stands for non-treated serving as a control group. (N = 5/treatment). Data show mean ± standard deviation (SD). Statistical significance was determined using a one-way ANOVA and Tukey’s multiple comparison post-test. *p < 0.05, ***p < 0.001, ****p < 0.0001.

### 3.3. rPS does not impair recovery following MRSA skin infection

Next, we asked whether rPS would worsen outcomes following infection. Hence, mice began rPS treatment at –7 dpi, were infected with MRSA midway through treatment, and were euthanized at 10 dpi (Fig. 3A). Rapamune PO 14x caused drastic weight loss lower than –10% relative to baseline (–7 dpi), highlighting a greater treatment burden after infection (Fig. 3B). In contrast, rPS mirrored the weight recovery of infected controls (MRSA) where both groups showed percent weight change above –10% throughout the monitoring period (Fig. 3B). Rapamune PO 14x produced significantly larger lesions than rPS at 3 and 5 dpi (Fig. 3C, D). Rapamune PO 14x was the only group with detectable bacteria in the liver (Fig. 3E). Bacterial burden at the skin lesion site was reduced for rPS-treated mice (Fig. 3F), supporting the good recovery outcomes observed in terms of weight and lesion area. Collectively, these findings indicate that rPS does not undermine the response necessary for rapid recovery following MRSA skin infection.

**Figure 3.**
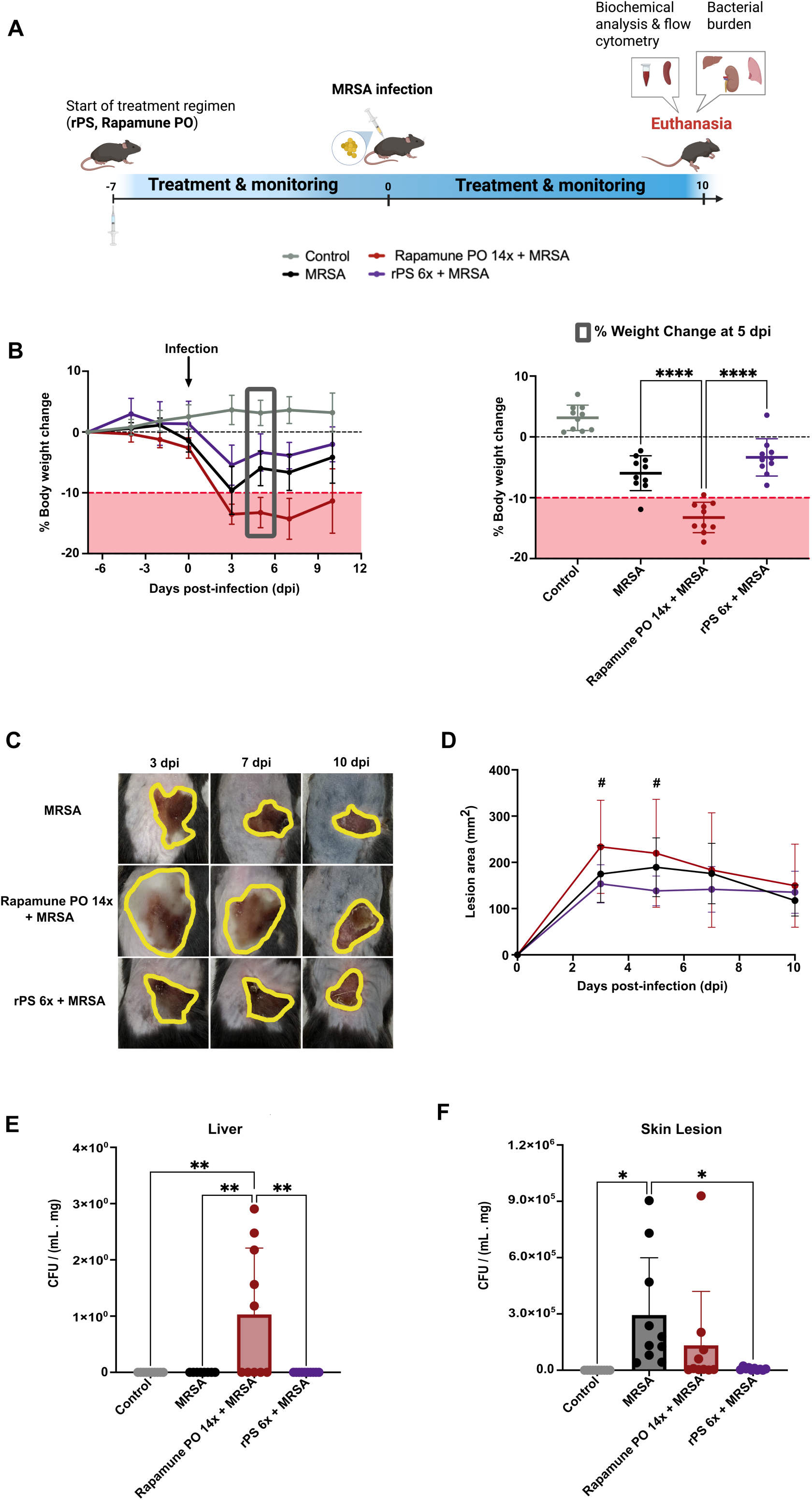
rPS maintains healthy body weight recovery following infection and prevents hepatic bacterial dissemination. (A) Experimental design of immunotherapy + MRSA infection study to evaluate the effect of rPS treatment on immune response to MRSA skin infection. Treatment with Rapamune PO 14x or rPS 6x began on –7 dpi and concluded on 7 or 8 dpi respectively. (B) Averaged Percent body weight change, calculated relative to body weight at the start of treatment (–7 dpi). The shaded area in red represents –10% to –20% weight loss which is considered a significant weight loss by the animal study protocol. The grey box represents the body weight change halfway through the infection monitoring period at 5 dpi and average percent change of body weight is compared between treatment groups. (C) Representative images of lesions induced following infection at highlighted timepoints. Yellow contours around the lesions represent the lesion areas. (D) Lesion area following infection in the dorsal region of the mice quantified using imageJ. # compares rPS 6x + MRSA vs Rapamune PO 14x + MRSA showing p < 0.05 at 3 and 5 dpi. (E) Bacterial burden in the liver 10 dpi normalized to the liver mass collected for each animal. (F) Bacterial burden in the lesion area of the skin at 10 dpi normalized to the mass of skin homogenized. (N = 10). Data show mean ± standard deviation (SD). Statistical significance was determined using a one or two-way ANOVA and Tukey’s multiple comparison post-test. *p < 0.05, **p < 0.01, ****p < 0.0001.

### 3.4. Distinct cytokine signatures in the serum and skin lesion

At 10 dpi, the skin lesion site harbored the greatest bacterial burden, suggesting that the bacteria and immune response would be primarily localized at the lesion site at this timepoint (Fig. 3F). Hence, we evaluated cytokines at the lesion and in the serum to compare inflammatory burden in the two environments (Fig. 4A). For all infected groups, the skin lesion exhibited a markedly more inflammatory environment, with TNF-□, IL-17A and IFN-γ significantly more enriched in the lesion relative to the serum than in the uninfected control (Fig. 4B, C, D). In contrast, IL-6 was not significantly elevated at 10 dpi at the lesion relative to the serum for infected rPS and Rapamune-treated mice compared to the control (Fig. 4E). Instead, we observed that increased IL-6 occurred earlier in the serum where all infected groups showed higher amounts of IL-6 than the control at 3 dpi (Supplementary Fig. S1).

**Figure 4.**
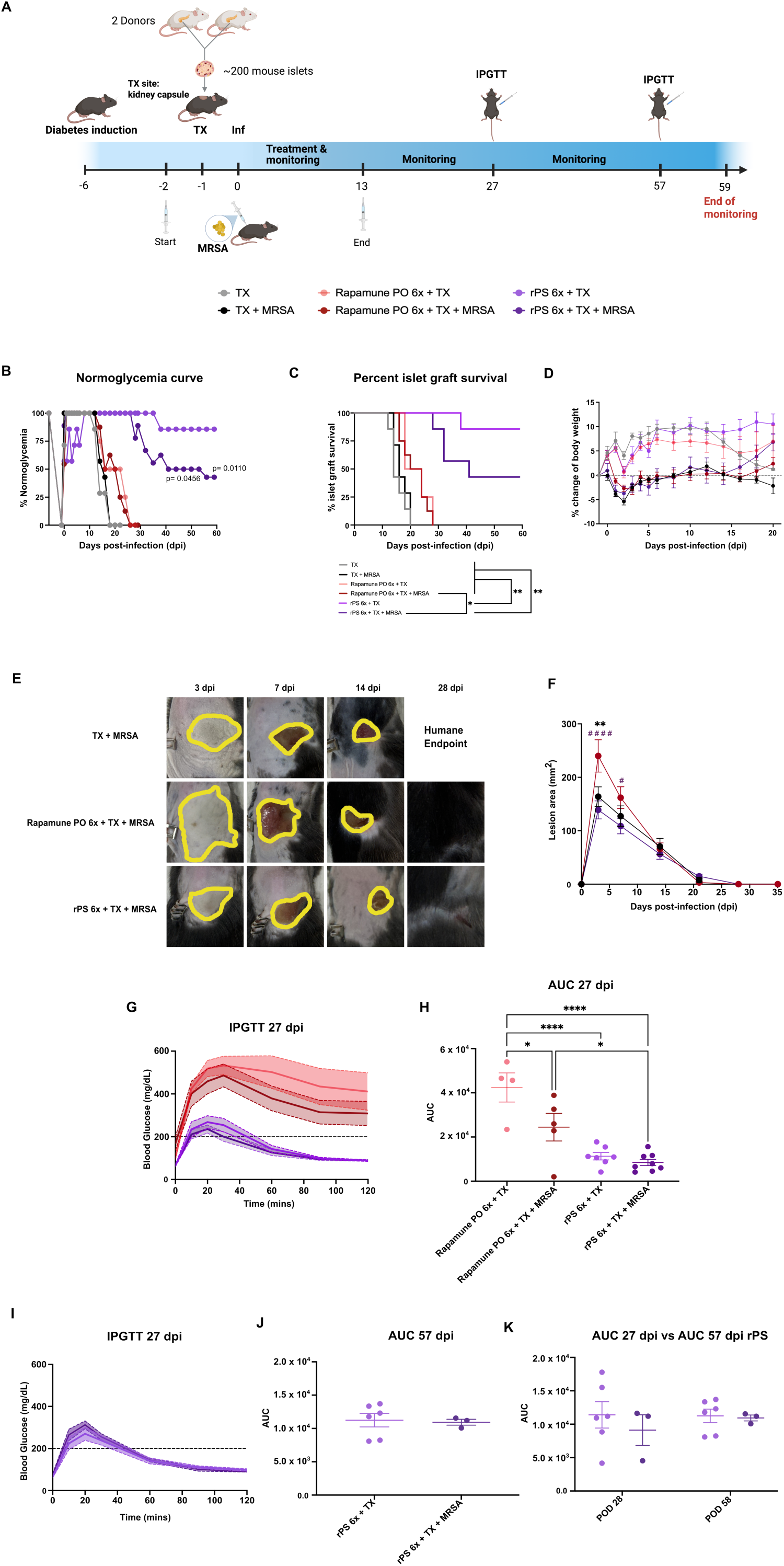
rPS promotes a Th2-skewed immune response to counteract MRSA driven localized inflammation at the lesion site. (A) Schematic summary of the experimental analysis performed in the serum, homogenized skin and spleen collected from mice euthanized at 10 dpi in the MRSA infection study. Log2 fold change comparing the levels of cytokines in the skin lesion and the serum at 10 dpi including (B) TNF-□, (C) IL-17A, (D) IFN-γ and (E) IL-6. (F) Flow Cytometry analysis of the frequency of IIL-17A+ CD4+ T cells (Th17 cells) in the spleen. (G) Frequency of CD11b+ Ly6G+ CD64+ neutrophils (Activated Neutrophils). Th2 response induced by rPS 6x shown by cytokine analysis in the serum and skin lesion shown via (H) IL-10 at the skin lesion, (I) IL-10 in the serum, (J) IL-4 at the skin lesion, and (K) IL-4 in the serum. (L) Increased frequency of IL4+ CD4+ splenic T cells (Th2 cells) induced by rPS leads to (M) decreased frequency of IFN-γ+ CD4+ splenic T cells (Th1 cells). N=10/treatment. Data show as mean ± SD (standard deviation). Statistical significance was determined using a one-way or two-way ANOVA and Tukey’s multiple comparison post-test. *p < 0.05, **p < 0.01, ***p < 0.001, ****p < 0.0001.

Analysis of splenic immune cells at 10 dpi revealed that T-helper 17 (Th17) cells were significantly reduced in the spleen across all infected groups, consistent with recruitment of Th17 cells from the spleen to the lesion site, where IL-17A levels were elevated (Fig. 4C, F). Activated neutrophils were similarly depleted from the spleen in infected groups and myeloperoxidase (MPO) activity was elevated at the lesion in Rapamune PO 14x-treated mice relative to rPS (Fig. 4G, Supplementary Fig. S2). Serum MPO levels did not differ among groups at 10 dpi (Supplementary Fig. S2). While rPS-treated mice showed signs of an inflamed lesion site, we observed reduced IFN-γ, IL-17A and MPO at the lesion compared to other infected groups (Supplementary Fig. S2, S4). Reduced inflammation in rPS-treated mice may reflect elevated IL-10 production at the lesion (Fig. 4H) and in serum (Fig. 4I). This anti-inflammatory response also included increased IL-4 at the lesion (Fig. 4J) and in the serum (Fig. 4K). This T-helper-2 (Th2) response was confirmed at the cellular level where rPS increased Th2 cell frequency (Fig. 4L) accompanied with a decrease in Th1 cells in the spleen (Fig. 4M). Collectively, these findings suggest that inflammation tilts towards the lesion rather than the systemic circulation at 10 dpi, and rPS induces a regulatory response that can promote inflammatory resolution without compromising pathogen control.

### 3.5. MRSA erodes long-term islet allograft survival induced by rPS

The prominent inflammation at the lesion site and the rPS-induced anti-inflammatory response prompted us to hypothesize that MRSA skin infection will not disrupt rPS-induced allograft survival given the distal nature of the inflammation relative to the graft location (kidney capsule). To test this hypothesis, STZ-induced diabetic C57BL/6 recipients of BALB/c islets were treated with rPS 6x or Rapamune PO 6x to allow a dose-matched comparison for graft survival. Recipients were then challenged subcutaneously with MRSA and monitored up to 60 days post-TX (Fig. 5A). Graft rejection was defined by two consecutive BG readings >300 mg/dL, whereas normoglycemia (BG <200 mg/dL) provided a graded measure of graft function when animals become hyperglycemic without meeting rejection criteria. Untreated TX recipients and TX + MRSA recipients were both universally hyperglycemic at 20 dpi (Fig. 5B, C, Supplementary Fig. S5), indicating that MRSA infection does not meaningfully accelerate rejection. Rapamune PO 6x did not significantly extend graft survival, and all Rapamune PO-treated recipients were hyperglycemic by 30 dpi (Fig. 5B, C, Supplementary Fig. S5). In contrast, rPS-treated recipients showed a clear infection-associated decline in graft function: 6/7 rPS 6x + TX animals remained normoglycemic through 59 dpi, compared with only 3/7 of infected recipients, with significantly lower normoglycemia rates in rPS 6x + TX at 50 (p = 0.0456) and 60 dpi (p = 0.0110) (Fig. 5B). The doubling of normoglycemia and survival rate (85.7% for rPS 6x + TX vs 42.8% for rPS 6x + TX + MRSA) at 59 dpi in uninfected animals demonstrates that MRSA skin infection measurably disrupted rPS-supported allograft acceptance (Fig. 5C).

**Figure 5.**
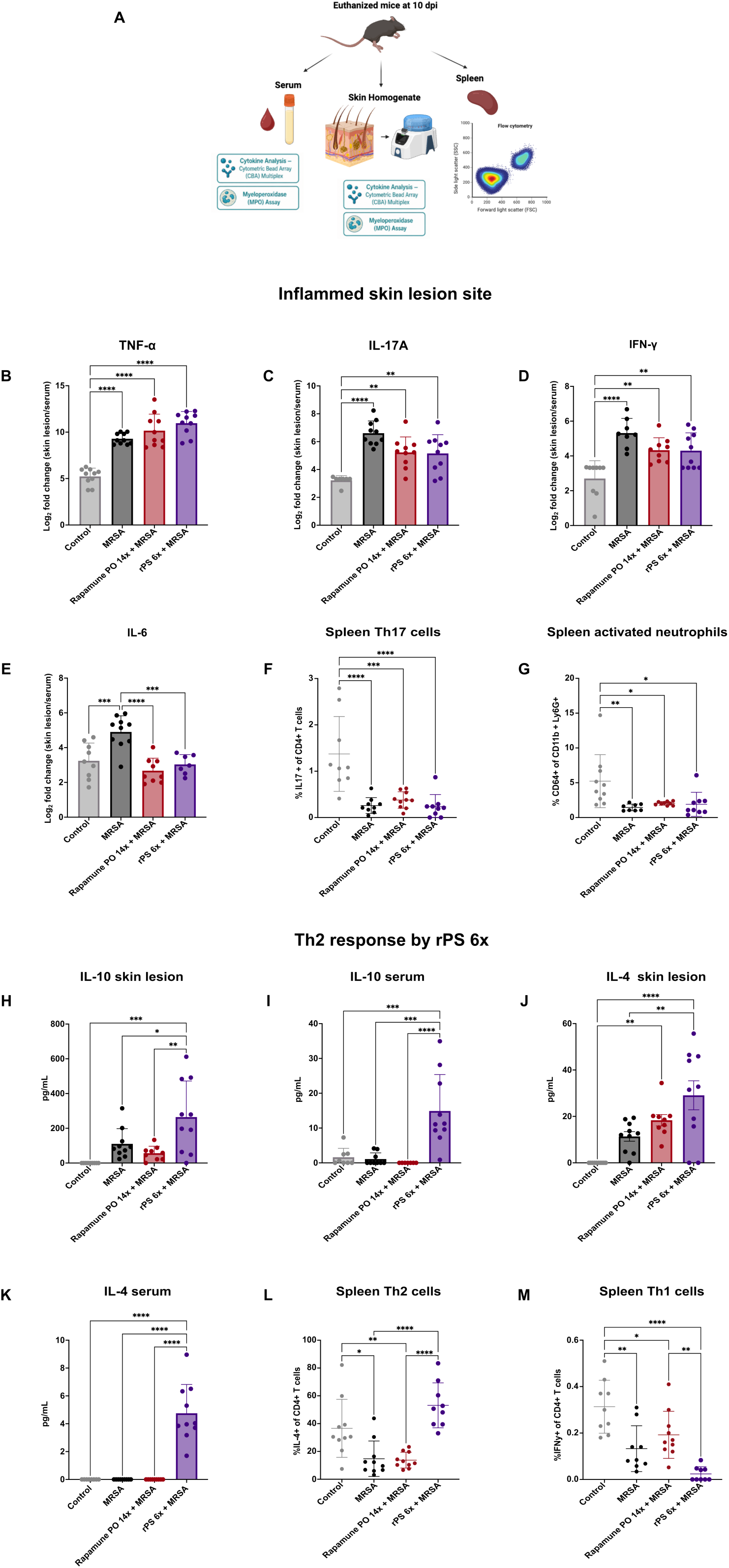
MRSA erodes superior allogeneic islet graft survival induced by rPS. (A) Experimental study design of immunotherapy + TX + MRSA infection. TX recipients received islets from 2 Balb/C donors in the kidney capsule and began treatment with rPS 6x or Rapamune PO 6x on –2 dpi. MRSA infection occurred 1 day after islet TX and recipients were monitored for up to 59 dpi. (B) Normoglycemia curve of TX recipients from diabetes induction on-6 dpi to 59 dpi. Normoglycemia is defined as blood glucose (BG) < 200 mg/dL. p-values compare rPS 6x + TX to rPS 6x + TX + MRSA on 50 and 59 dpi. (C) Survival curve depicting protection of the transplanted islet graft. Graft failure is defined as two consecutive BG > 300 mg/dL. (D) Averaged percent change of body weight, calculated relative to weight on TX day (–1 dpi) up to 20 dpi. (E) Representative lesion images of infected mice at 3, 7, 14 and 28 dpi. No mice in the TX + MRSA group were non-diabetic by 28 dpi and were humanely euthanized. (F) Quantified lesion area following MRSA infection in the dorsal region of TX recipients. ** compares TX + MRSA vs Rapamune PO 6x + TX + MRSA showing p < 0.01. #### and # compares rPS 6x + TX + MRSA vs Rapamune PO 6x + TX + MRSA showing p < 0.0001 and p < 0.05, respectively. (G) Intraperitoneal glucose tolerance test (IPGTT) administered to TX recipients with surviving grafts on 27 dpi. Solid lines show average values and shading corresponds to SEM. (H) IGPTT area under the curve (AUC) on 27 dpi. (I) IPGTT on 57 dpi for surviving recipients in rPS 6x + TX and rPS 6x + TX + MRSA. (J) AUC on 57 dpi. (K) Comparison of AUC on 28 vs 58 dpi of TX recipients in rPS 6x + TX and rPS 6x + TX + MRSA with surviving grafts at both timepoints. (N = 4-9). Data show mean ± SEM. Survival curves were compared using a Log Rank test and Holm-Šídák post-hoc analysis for multiple comparisons. Statistical significance was determined using a one-way or two-way ANOVA and Tukey’s multiple comparison post-test. *p < 0.05, ****p < 0.0001.

As for post-infection recovery, all infected groups showed transient weight loss after infection (Fig. 5D). rPS-treated recipients returned to ∼0% by 14 dpi and steadily gained weight thereafter, while untreated and Rapamune PO-treated recipients trended toward weight loss as hyperglycemia progressed (Fig. 5D, Supplementary Fig. S6). Rapamune PO 6x produced the largest lesions, exceeding TX + MRSA (p < 0.01, 3 dpi) and rPS 6x + TX + MRSA (p < 0.0001 at 3 dpi, p < 0.05 at 5 dpi), though lesions for all treated groups were closed by 35 dpi (Fig. 5E, F). IPGTT at 27 dpi revealed superior glucose clearance for rPS recipients relative to Rapamune PO recipients, whose glucose curves peaked near 600 mg/dL with correspondingly larger areas under the curve (AUC) (Fig. 5G, H). By 57 dpi, when only rPS-treated recipients remained, infected and uninfected mice showed equivalent glucose clearance (Fig. 5I, J), a pattern also observed at 27 dpi. Restricting analysis to rPS-treated recipients surviving at both 27 and 57 dpi revealed no difference in glucose clearance between infected and uninfected animals, highlighting that glucose clearance is preserved in survivors regardless of infection status (Fig. 5K). Collectively, these data show that MRSA skin infection erodes rPS-induced allograft acceptance, yet surviving infected recipients retain glucose-stimulated insulin responsiveness similar to uninfected rPS-treated TX recipients.

### 3.6. Early burst of IL-6 later followed by TNF-□ and neutrophilia drive MRSA-induced erosion

To elucidate the mechanisms promoting MRSA erosion of rPS-induced tolerance, we performed cytokine and complete blood count (CBC) analyses in the serum and whole blood respectively (Fig. 6A). All infected groups showed a significant increase in IL-6 relative to their uninfected controls at 16 hpi (p < 0.0001) and 1 dpi (p < 0.01), an increase that was transient, declining sharply between these two timepoints and declining back to nearly undetectable levels by 13 dpi (Fig. 6B). Average TNF-levels, in contrast, rose progressively among rPS– and Rapamune PO-treated recipients through 29 dpi (Fig. 6C). Infected rPS 6x + TX + MRSA recipients showed significantly higher TNF-than TX + MRSA and all uninfected controls at 13 dpi (Fig. 6C). These inflammatory signals likely overrode regulatory cytokines in their ability to erode tolerance as IL-10 and IL-4 modestly increased at 13 and 29 dpi for rPS-infected recipients (Supplementary Fig. S7).

**Figure 6.**
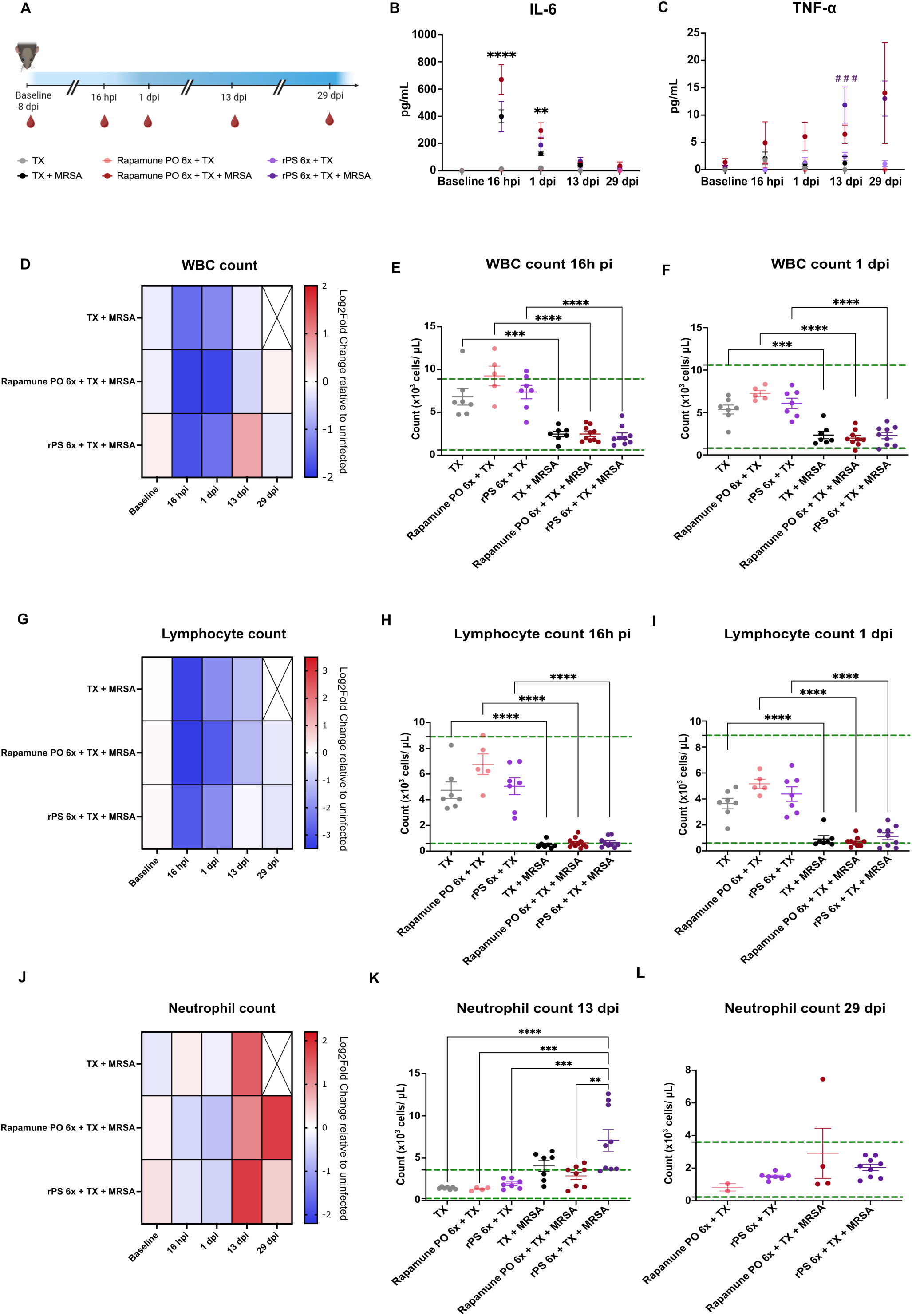
Early burst of IL-6 later followed by neutrophilia drives MRSA erosion of islet graft survival induced by rPS 6x. (A) Schematic of retro-orbital bleeding timepoints of TX recipients at baseline (–8 dpi) 16 hpi, 1dpi, 13 dpi and 29 dpi for cytokine and complete blood count (CBC) analyses. (B) IL-6 levels in serum with significance comparison between infected and the uninfected counterparts at 16 hpi and 1 dpi (i.e. TX + MRSA vs MRSA ****p <0.0001). (C) TNF-□ levels in serum with significance comparison # # # # p < 0.0001 between rPS 6x + TX + MRSA and all uninfected cohorts (TX, Rapamune PO 6X + TX and rPS 6x + TX). (D) Categorical heatmap showing Log2 fold change of white blood cell (WBC) count of an infected group relative to their corresponding uninfected group (i.e., rPS 6x + TX + MRSA relative to rPS 6x. + TX). (F) WBC count at 16 hpi and (F) 1 dpi comparing all TX recipients. (G) Heatmap of Log2 fold change of lymphocyte count at indicated timepoints. (H) Lymphocyte count at 16 hpi and (I) 1 dpi. (J) Neutrophil count heatmap of Log2 fold change relative to uninfected cohort. (K) Neutrophil count at 13 dpi and (L) 29 dpi. Dashed green lines represent the reference ranges of WBC, lymphocyte, and neutrophil counts for healthy wild-type mice provided by the instrument. (N = 4-9). Data show mean ± SEM. Statistical significance was determined using a one-way or two-way ANOVA and Tukey’s multiple comparison post-test. **p < 0.01, ***p < 0.001, ****p < 0.0001.

All infected groups showed a decline in average white blood cell (WBC) count at 16 hpi and 1 dpi relative to uninfected controls (Fig 6D). The WBC counts among all infected recipients fell closer to the leukopenic threshold of 0.8 x 10³ cells/μL, while uninfected recipients remained within the reference range of 0.8-10.6 x 10³ cells/μL (Fig. 6D-F). This WBC decline was driven by significant reduction in lymphocyte counts. Indeed, all infected recipients showed average lymphocyte counts below the lymphopenic threshold (<0.6 x 10³ cells/μL) at 16 hpi (Fig. 6G, H). Lymphocyte counts remained significantly lower in infected versus uninfected recipients at 1 dpi (p < 0.001, p < 0.0001). However, average values in TX + MRSA and rPS-treated infected recipients had recovered above the lymphopenic threshold by 1 dpi, confirming the transient nature of the lymphopenia (Fig. 6I).

Neutrophil counts, by contrast, barely changed initially after infection (Fig. 6J). They then rose sharply at 13 dpi across all infected groups relative to uninfected controls (Fig. 6J, K). Infected rPS-treated recipients showed the most pronounced neutrophilia (Fig. 6K). Furthermore, monocyte counts also increased at 13 dpi among infected groups while remaining within their reference range (Supplementary Fig. S8). As with the other infection-associated hematologic changes, the neutrophilia was not sustained. Average neutrophil counts in most surviving infected recipients returned to the reference range by 29 dpi (Fig. 6L). Collectively, these data indicate that MRSA infection triggers an early, transient IL-6 burst accompanied by significant lymphocyte depletion from the circulation within 24h of the infection. A subsequent wave of neutrophilia and TNF-at 13 dpi collectively represent sequential inflammatory insults that can disrupt rPS-induced graft survival.

### 3.7. Combination of rPS with methylprednisolone (MP) reduces IL-6 and increases islet survival despite MRSA infection

Given previous reports identifying IL-6 as a major barrier to tolerance induction in skin allografts,^14,30^ we speculated that blunting IL-6 production which can also lower TNF-levels,^31,32^ will permit increased graft survival among infected rPS-treated TX recipients. As such, we combined rPS with methylprednisolone (MP), a glucocorticoid used in TX induction protocols to lower inflammation associated with the administration of lymphocyte-depleting agents.^33,34^ Hence, a single 20 mg/kg dose of MP was administered on the day of infection with all other parameters unchanged (Fig. 7A). The combination therapy increased normoglycemia and graft survival to 80% in infected recipients, nearly doubling the rate achieved among infected rPS recipients (42.8%) (Fig. 7B, C, Supplementary Fig. S9). Without rPS and presence of MP alone, all recipients became diabetic by 20 dpi, confirming that presence of rPS is required for tolerance induction (Fig. 7B, C). Uninfected recipients receiving either rPS alone or the combination therapy reached 85.7% and 80% graft survival rate respectively, indicating that MP shields rPS-induced tolerance from MRSA-driven erosion (Fig. 7B, C). Mechanistically, the graft survival benefit of combination therapy tracked with a significant reduction in IL-6 at 16h post-infection relative to rPS alone (Fig. 7D). TNF-was elevated only in infected rPS-treated recipients, revealing that lowering IL-6 also lead to decreased TNF-among infected recipients treated with rPS + MP (Figs. 6C, 7E). Lymphopenia at 16 hpi and neutrophilia at 13 dpi persisted in both rPS alone and rPS + MP infected groups, indicating that restoring superior allograft acceptance did not require downregulation of all inflammatory insults (Fig. 7H, I). Together, these results show that addition of MP to the rPS treatment regimen selectively dampened early IL-6 and subsequent TNF-production, restoring graft survival to levels comparable to uninfected TX recipients.

**Figure 7.**
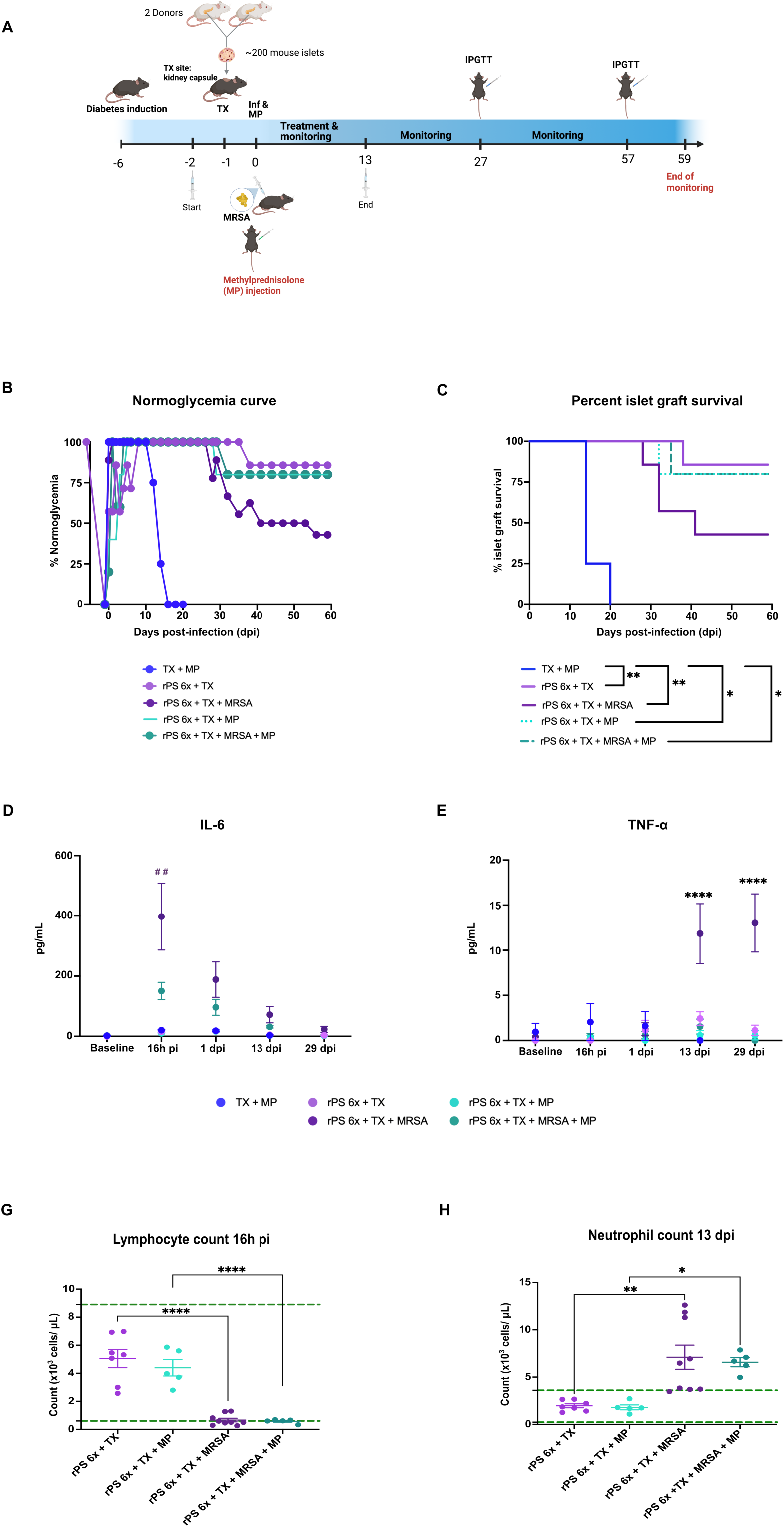
Combination between rPS and methylprednisolone (MP) reduces MRSA-induced IL-6 and restores higher islet allograft acceptance. (A) Experimental study design following the same design as Fig. 5A except for the administration of MP on the day of MRSA infection. (B) Normoglycemia curve of TX recipients from diabetes induction on – 6 dpi to 59 dpi. rPS 6x + TX and rPS 6x + TX + MRSA are the same as shown in Fig. 5. Normoglycemia is defined as blood glucose (BG) < 200 mg/dL. (C) Allogenic islet graft survival up to 60 days post-TX. Graft failure is defined as two consecutive BG > 300 mg/dL. (D) IL-6 levels in the serum at the indicated timepoints after infection. # # compares rPS 6x + TX + MRSA vs rPS 6x + TX + MRSA + MP showing p < 0.01. (E) TNF-□ levels in the serum with **** showing comparisons between rPS 6x + TX + MRSA and any other treatment group at 13 and 29 dpi (i.e., rPS 6x + TX + MRSA vs TX + MP). (F) Lymphocyte count at 16 hpi. (G) Neutrophil count at 13 dpi. Dashed green lines represent the reference range for lymphocyte and neutrophil counts. (N = 4-9). Data show mean ± SEM. Statistical significance was determined using a one-way or two-way ANOVA and Tukey’s multiple comparison post-test. *p < 0.05, **p < 0.01, ****p < 0.0001.

## Discussion

This work examined whether a targeted nanotherapy for islet TX would weaken host defense against MRSA skin infection, and whether the resulting antibacterial response from a distant infection relative to graft location can erode islet tolerance established by the nanotherapy. Our findings support an important difference between antimicrobial protection and the durability of allograft survival. At the cellular level, rPS inhibited ROS production and NETosis, yet unlike rapamycin, rPS preserved bactericidal capacity and neutrophil viability. These results align with work by Genito et al. showing that mTOR signaling is necessary for free radical production of neutrophils.^24^ *In vivo*, rPS-treated mice rapidly recovered from the subcutaneous MRSA infection without signs of disseminated bacteria in the liver and low bacterial burden at the lesion. These results are consistent with the ability of rPS to deliver rapamycin to antigen-presenting cells in lymph nodes away from circulating immune cells that can be recruited to the site of infection.^19^

MRSA is the leading cause of skin and soft-tissue infections which are frequent infection complications among transplant recipients in the hospital setting.^35,36^ Despite a preservation of antimicrobial defenses, cutaneous MRSA infection still disrupted rPS-induced tolerance. In this skin infection model, the infection produced a compartmentalized inflammatory signature, with TNF-□, IL-17A, and IFN-γ enriched at the inoculation site. This localized immune signature was also reported by Chan et al. where pro-inflammatory cytokines were more prominent in the skin than plasma and spleen.^37^ Furthermore, the decreases in WBC and lymphocyte counts in the blood observed among infected TX recipients is consistent with accumulation of inflammatory cells at the site of infection as previously reported.^38^ Amid this localized inflammation, rPS induced a notable Th2-skewed response that has been associated both with improved resolution of bacterial infection and worsening outcomes as the bacteria can upregulate regulatory mechanisms to evade the immune response.^39–41^ Given that rPS-treated mice showed reduced lesion bacterial burden, we posited that this response served to resolve the inflammation. Still, the localized inflammation and regulatory Th2 response induced by rPS at 10 dpi had little bearing on preventing erosion of rPS-induced graft survival despite the distal infection relative to graft location. This discrepancy is best explained by the timing and environment in which inflammation was evaluated. Although bacteria was largely concentrated at the lesion at 10 dpi, the early response was systemic as shown by the rapid IL-6 serum increase used to activate T cells for neutrophil recruitment to the site of infection.^42^ Because the lesion site was not harvested during the first 24h, the initial contribution of localized and systemic responses cannot be distinguished in the present study. Therefore, the data support the conclusion that a distal MRSA lesion generated an early systemic inflammatory relay capable of perturbing allograft-induced survival. These findings also highlight the importance of infection routes in this infection-breaking tolerance model. Systemic versus pulmonary MRSA infection would generate cytokine levels across different microenvironments and timepoints, underscoring the difficulty of generalizing these results to other MRSA infection models.

Accordingly, our findings are most informative when tolerance is understood as a graded continuum of robustness rather than as a binary outcome.^17,43^ Indeed, MRSA did not eliminate rPS-mediated graft protection in every recipient; instead it lowered the proportion that maintained long-term graft survival. Consistent with this graded continuum of tolerance, a single dose of MP reduced early IL-6 and subsequent TNF-elevation without modifying the neutrophilia. Hence, the identity, timing, and magnitude of particular inflammatory signals appeared more consequential than the total inflammatory cascades as tolerance is induced by overlapping mechanisms that may have different susceptibilities to distinct inflammatory insults.^15^ Furthermore, we cannot exclude the possibility that IL-6 and TNF-□ worked synergistically to erode allograft acceptance.^30^

A limitation of this work is that we did not test the depth of the eroded tolerant state using immune-supercharging agents such as anti-PD-L1, anti-CD25 or future transplantability in infected recipients with failed grafts. Prior cardiac allograft work shows that grafts surviving a *Listeria monocytogenes* infection-induced rejection crisis can harbor a checkpoint-dependent tolerance that appears stable but is functionally fragile, since anti-PD-L1 alone precipitates rejection in these recipients but not in uninfected controls.^15^ While the surviving rPS-treated, MRSA-infected recipients retained glucose clearance comparable to rPS 6x + TX, we cannot rule out that these grafts are being maintained by a similarly eroded state rather than by a fully restored tolerogenic signature. Additionally, the possibility of a second transplant in infection-driven rejected recipients warrants further study as infection with *Listeria monocytogenes* has shown to allow for spontaneous restoration of tolerance in a second transplant whereas infection with cytomegalovirus showed accelerated rejection in a second transplant.^17,18^ These opposing findings highlight the importance of examining the pathogen-specific characteristics, the transplanted organ and tolerance induction strategies.

In conclusion, rPS can preserve effective neutrophil antibacterial activity while the induced tolerance remains vulnerable to an early systemic inflammation generated by a distal cutaneous infection relative to graft location. Finally, our findings also demonstrate that graft acceptance can be restored without eliminating all infection-associated inflammation.

## Author contributions

E.H.A.M. synthesized and characterized rPS and blank PS and performed all *in vivo* experiments presented in the manuscript. C.P.R. performed neutrophil experiments and A.J.M. supervised and assisted in the design of *in vitro* experiments. D.B.S., S.Z., N.R.K., S.A., A.B.K.B., R.R.B.M. and N.M. assisted in all *in vivo* experiments. X.Z. performed islet isolations and transplants in the kidney capsule. S.A. assisted in flow cytometry analysis and SAXS data collection and analysis. E.H.A.M, D.B.S., E.A.S. and J.A.B designed and conceived all experiments. E.H.A.M., D.B.S., E.A.S and J.A.B wrote and edited the manuscript. All authors read and approved of the final version of the manuscript.

## Data availability

Any additional data related to this work is available from the corresponding author upon reasonable request.

## Acknowledgements

This work made use of the Keck-II facility (RRID: SCR_026360) of Northwestern University’s NUANCE Center, which secured support from the International Institute of Nanotechnology and Northwestern University’s MRSEC program (NSF DMR-2308691). SAXS experiments were performed at the DuPont-Northwestern-Dow Collaborative Access Team (DND-CAT) which received funding from Northwestern University, The Dow Chemical Company, and DuPont de Nemours, Inc. Cryo-TEM images were recorded at the University of Virginia Molecular Electron Microscopy Core facility (RRID: SCR_019031), which receives support in part by the School of Medicine. The authors acknowledge the Center for Comparative Medicine Pre-Clinical Services for assistance with running the Element HT5 hematology analyzer for complete blood count analysis

## Data availability

No high-complexity data sets were used in the preparation of this manuscript.

## Supplementary data

All supplementary data to this manuscript can be found in the attached supporting information document.

## Generative AI statement

No generative AI or large learning model was used for data analysis and scientific interpretation. Perplexity was used for standard grammar to refine language and improve readability of the manuscript. The final version of the manuscript was reviewed by all authors and the content was edited as deemed necessary. All authors take full responsibility for the content of this publication.

## Abbreviations

AUC: area under the curve
BG: blood glucose
CBC: complete blood count
CFU: colony-forming unit
dpi: day post-infection
hpi: hours post-infection
IL-6: interleukin-6
IL-4: interleukin-4
IL-10: interleukin-10
IFN-γ: interferon-gamma
IPGTT: intraperitoneal glucose tolerance test
MRSA: Methicillin-resistant *Staphylococcus aureus*
MP: methylprednisolone
NT: non-treated
rPS: rapamycin-loaded polymersomes
Stim: stimulated
TX: transplantation;
TNF-□: tumor necrosis factor alpha
Th1: T-helper 1
Th2: T-helper 2
Th17: T-helper 17

