## Supplementary Information for "Preserving Nanotherapy-Induced Allogeneic Murine Islet Graft Tolerance During Distal Methicillin-Resistant Staphylococcus aureus Skin Infection"

**Title:**

Débora B. Scariot^1,3^

Xiaomin Zhang^4^

Calleigh P. Reber^5^

Natalie R. Klug^1,2^

Szumo Wang^1,2^

Sultan Almunif^1,6^

Amanda B. K. Bakoshi^1^

Rayanne R. M. Machado^1^

Natalia Mendoca^1^

Andrew J. Monteith^5^

Evan A. Scott^1,3^

Jacqueline A. Burke^1,2^

**Affiliations:**

1. Department of Biomedical Engineering, Northwestern University, Evanston, IL 60208, USA
2. Querrey Simpson Institute for Regenerative Engineering, Northwestern University, Chicago, IL 60611, USA
3. Department of Biomedical Engineering, NanoSTAR Institute, University of Virginia School of Medicine, Charlottesville, VA 22904, USA
4. Comprehensive Transplant Center, Feinberg School of Medicine, Northwestern University, Chicago, IL, USA
5. Department of Microbiology, University of Tennessee, Knoxville, TN 37996, USA
6. Bioengineering Institute, King Abdulaziz City for Science and Technology, Riyadh 11442, Saudi Arabia

**Address correspondence to:**

Jacqueline Burke, PhD

Querrey Simpson Institute for Regenerative Engineering, Northwestern University

Robert H. Lurie Medical Research Center, Room 11-111

Chicago, IL 60611

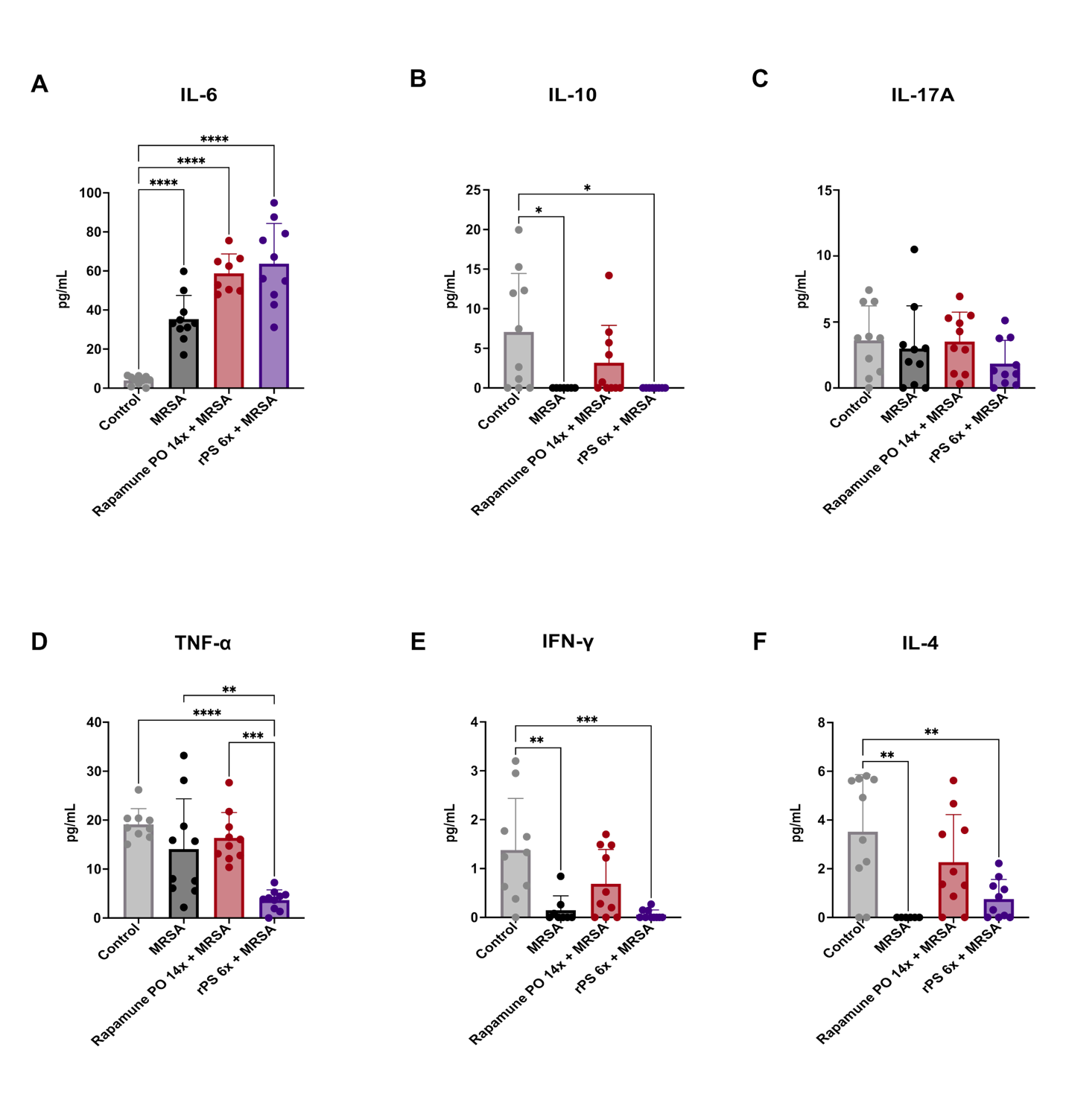

**Supplementary Figure S1: Cytokine analysis in serum at 3 dpi in immunotherapy + MRSA infection study.** Cytokine levels in serum following retro-orbital bleeding at 3 dpi measuring **(A)** IL-6 elevated for all infected groups, **(B)** IL-10, **(C)** IL-17A, **(D)** TNF-ɑ**, (E)** IFN-γ, **(F)** IL-4. Data show mean ± standard deviation (SD).

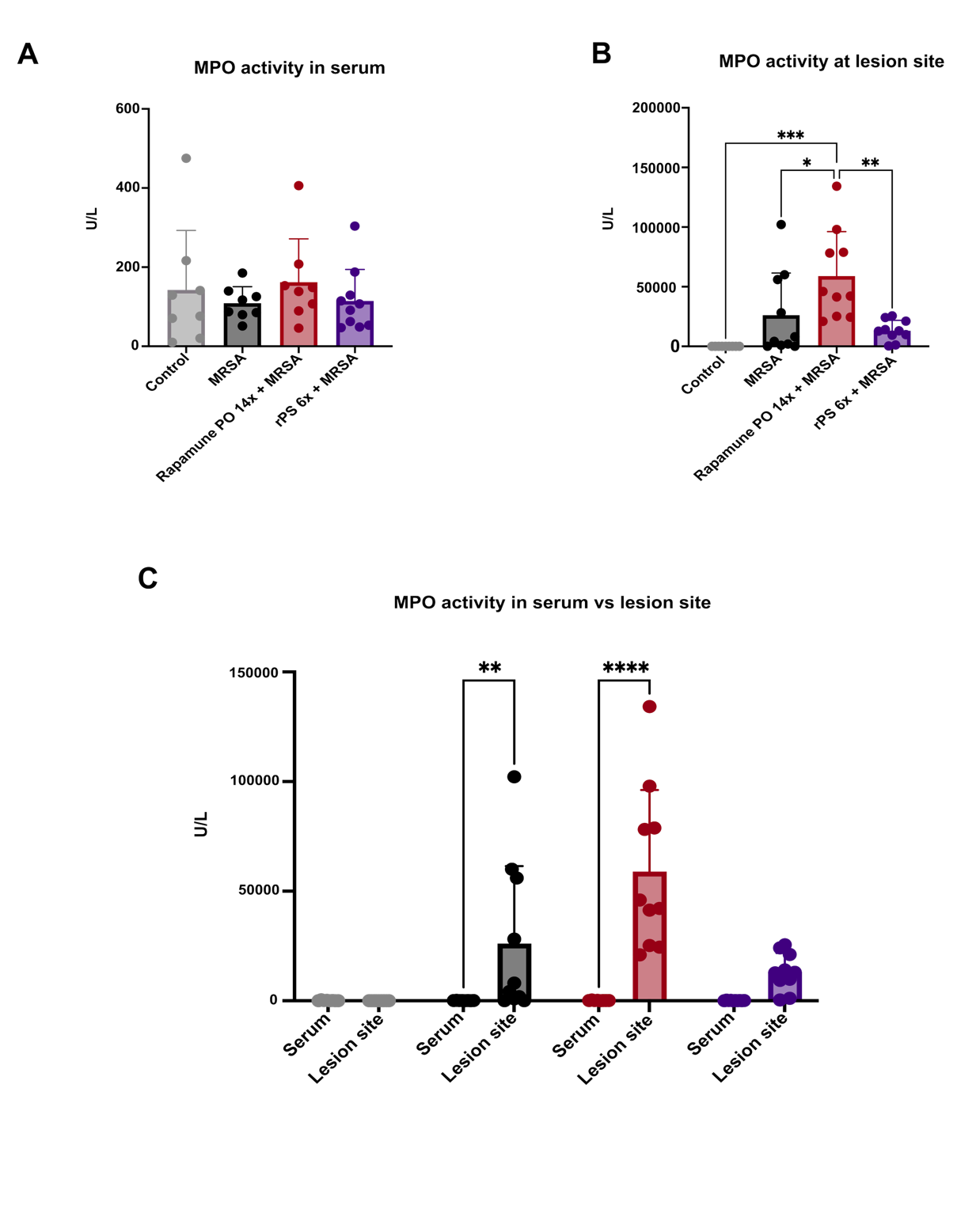

**Supplementary Figure S2: Myeloperoxidase (MPO) levels in serum and at the lesion site at euthanasia or 10 dpi. (A)** Comparable levels of MPO at 10 dpi in the serum for all treatment groups. **(B)** MPO levels at the lesion site with tissue mass corrections showing elevated MPO levels for rPS 14x. **(C)** MPO levels comparison between serum and lesion site. Data show mean ± standard deviation (SD).

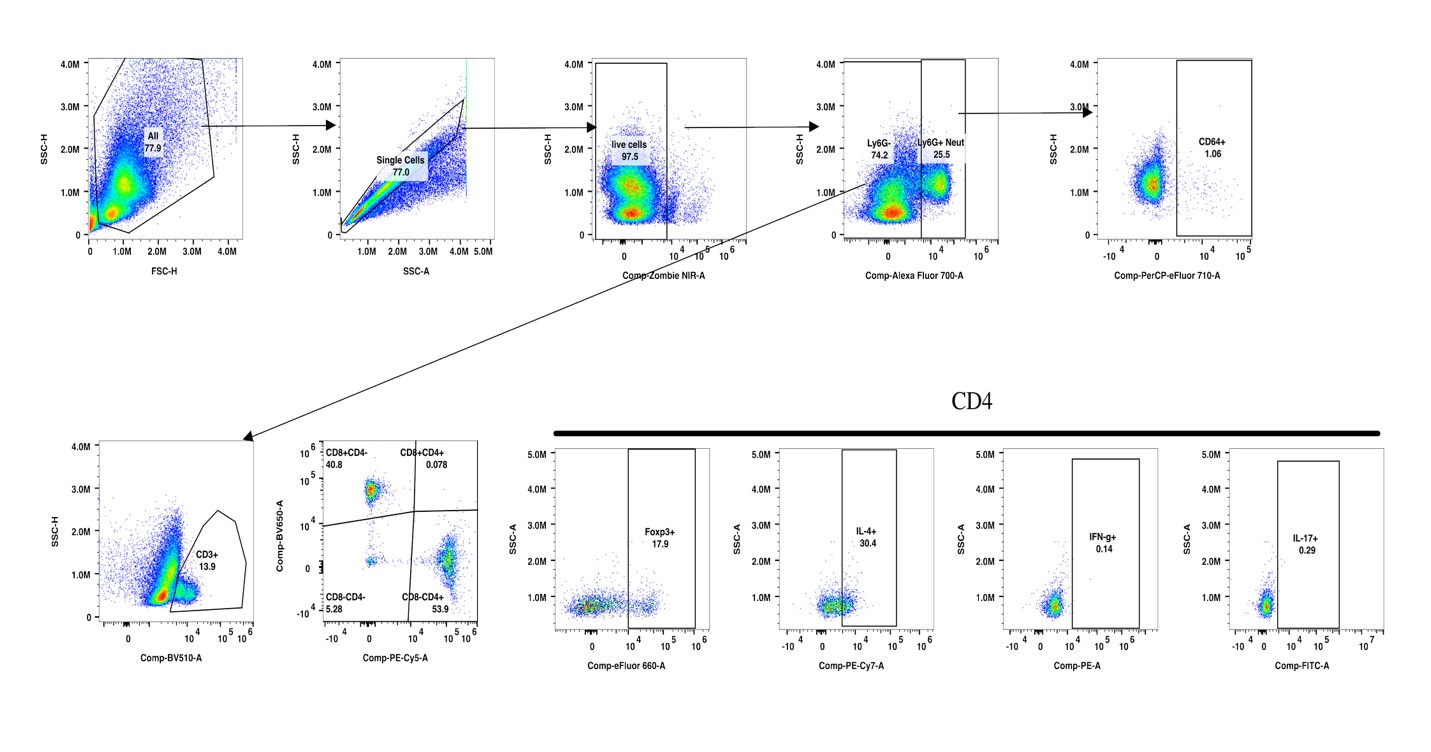

**Supplementary figure S3: Gating strategy for flow cytometry of the spleen in immunotherapy + MRSA infection study.** From all recorded events to singlets, live cells, activated neutrophils (Ly6G+ CD64+), and CD8-CD4+ T helper cells.

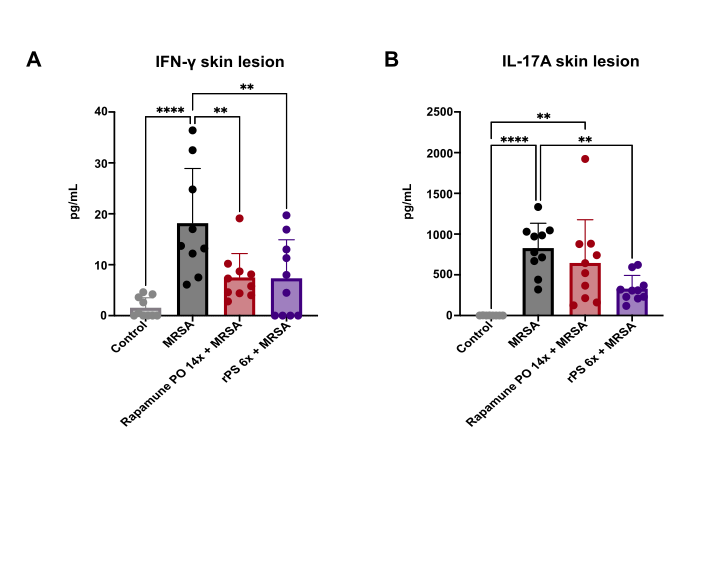

**Supplementary Figure S4:** rPS treated mice show lower IL-17 and IFN-γ at the lesion site compared to other infected groups. **(A)** IFN-γ, **(A)** IL-17 Data show mean ± standard deviation (SD).

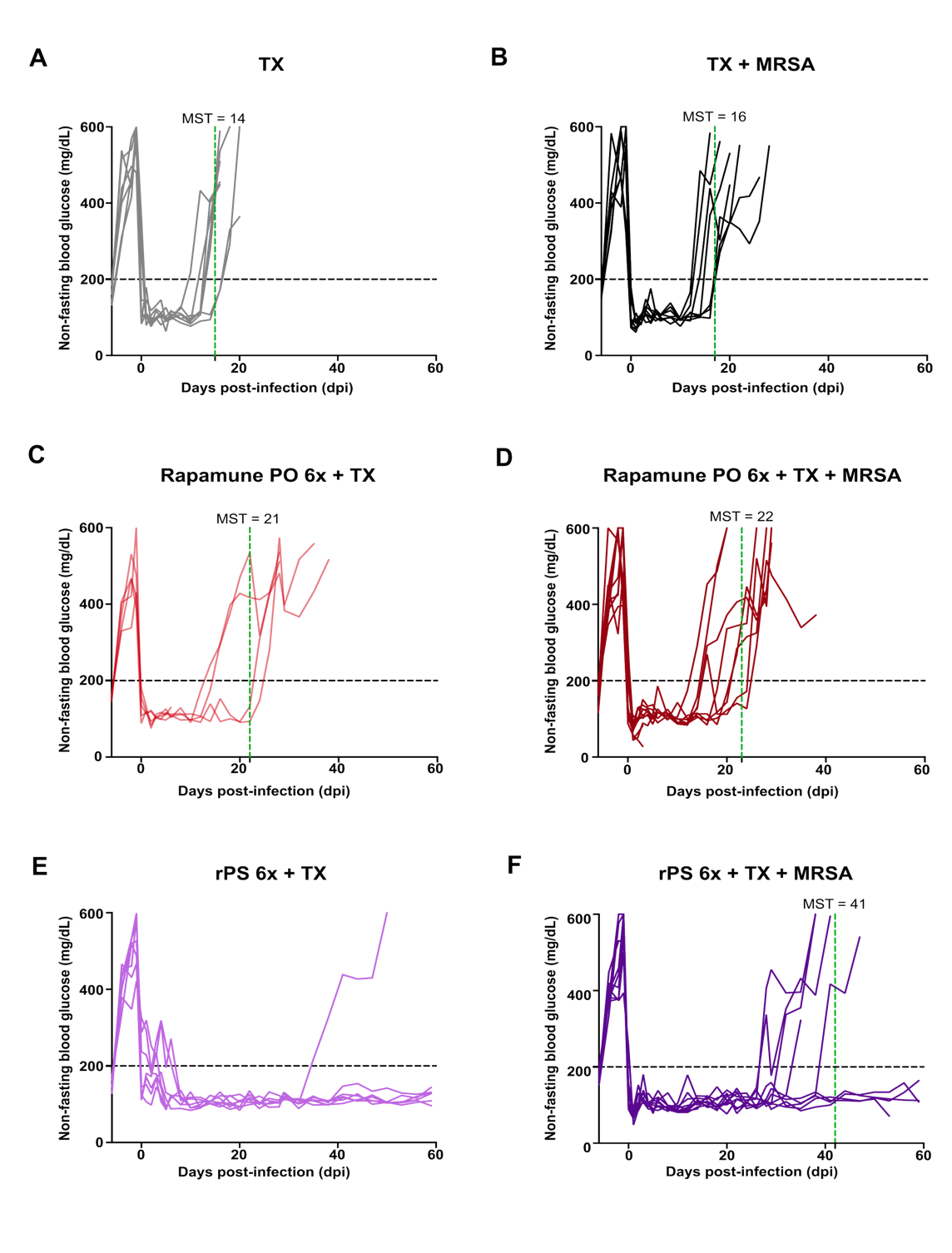

**Supplementary Figure S5: Blood glucose (BG) levels from -6 dpi (Streptozotocin injection day) to 59 dpi. (A)** TX (N=7) with median survival time (MST) of 14 days defined as the length of time when half of the transplant recipients have rejected islet graft with two consective BG > 300 mg/dL. **(B)** TX +MRSA (N=7) with MST =16 days. **(C)** Rapamume PO 6X + TX (N=5) with MST = 21 days. One recipient died unrelated to high BG levels. **(D)** Rapamune PO 6x + TX + MRSA (N=11) with MST = 22 days. Three recipients died unrelated to high BG levels. Death was attributed to weakness following Rapamune PO treatment and MRSA infection. **(E)** rPS 6x + TX (N=7) with undefined MST. **(F)** rPS 6x + TX + MRSA (N=9) with MST of 42 days. Two recipients died unrelated to high BG levels. Death was attributed to weakness after anesthesia with isoflurane.

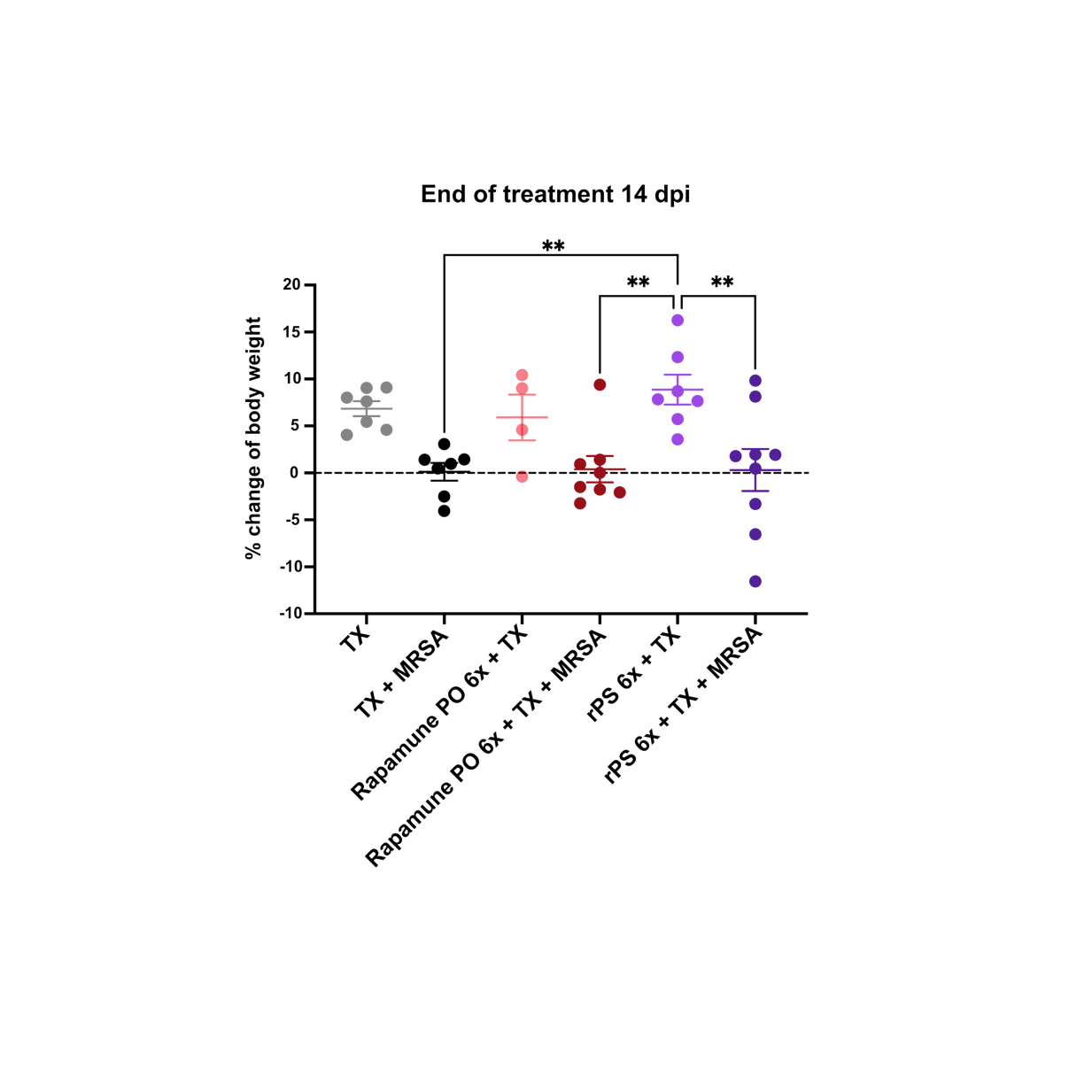

**Supplementary Figure S6: Percent change of body weight at 14 dpi with treatment beginning on -2 dpi and ending on 13 dpi.** Averaged percent change of body weight based on weight on day of transplantation at -1 dpi demonstrating MRSA infection-induced body weight loss for infected groups. Data show mean ± standard error of the mean (SEM).

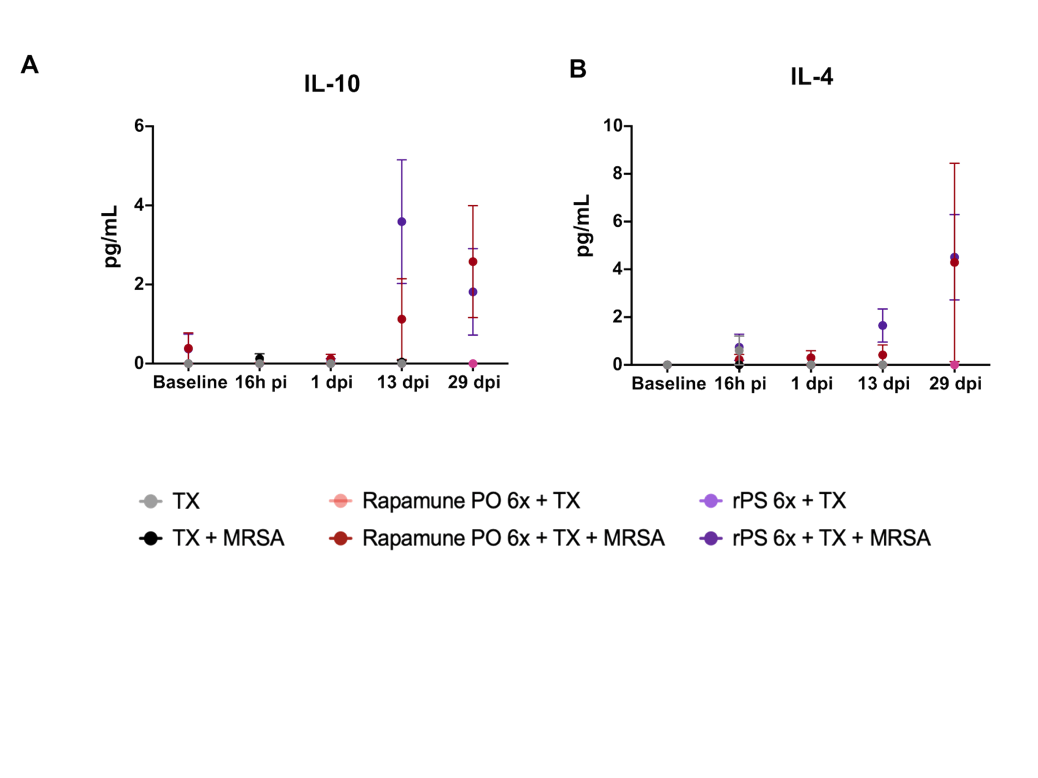

**Supplemental Figure S7: Cytokine levels in the serum of transplant recipients at specified timepoints.** Increased levels of **(A)** IL-10 and **(B)** IL-4 rPS-infected recipients despite TX erosion.

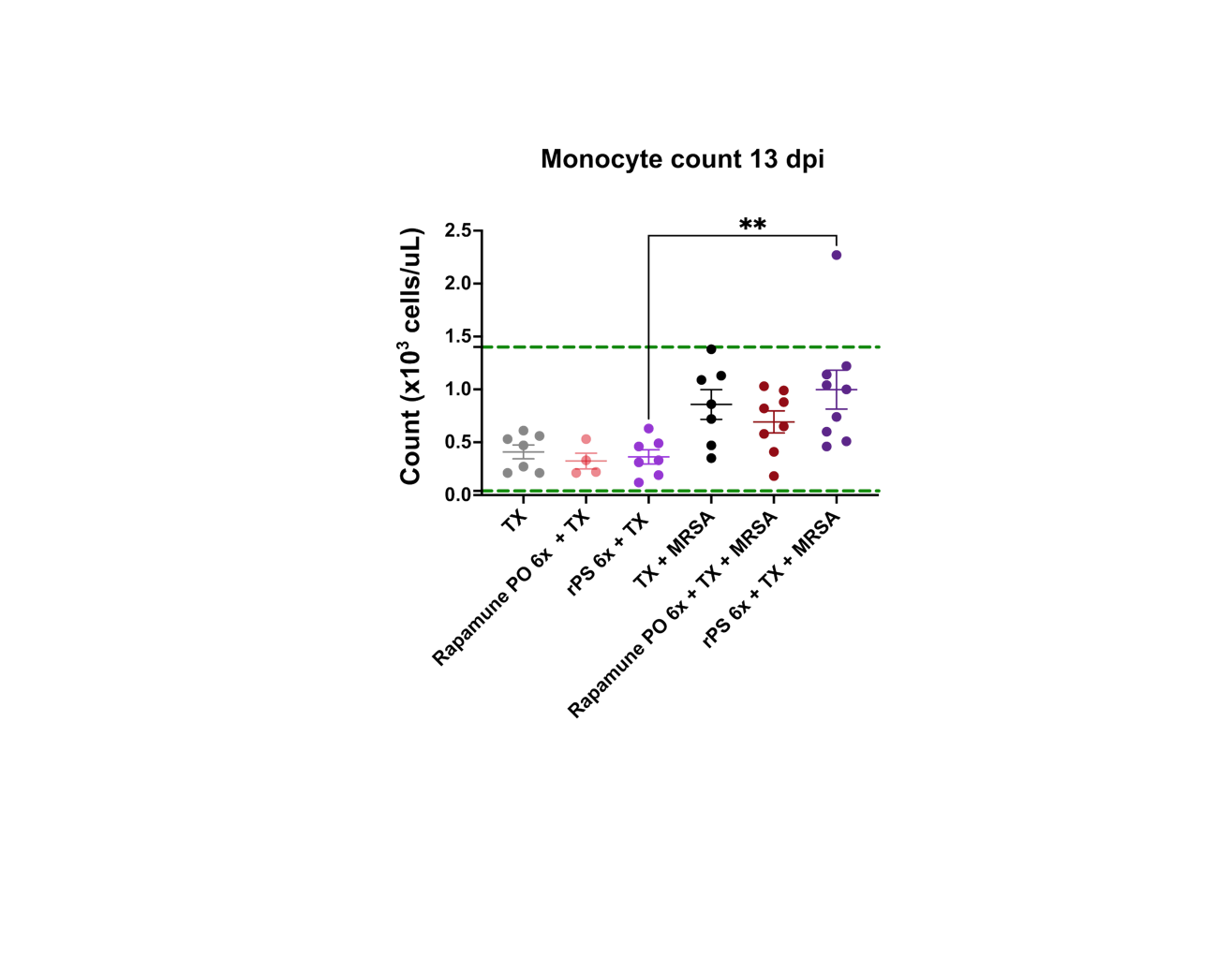

**Supplementary Figure S8: Increase in monocyte counts for infected groups, particularly for rPS-treated recipients.** Dashed green lines represent the reference range for monocytes.

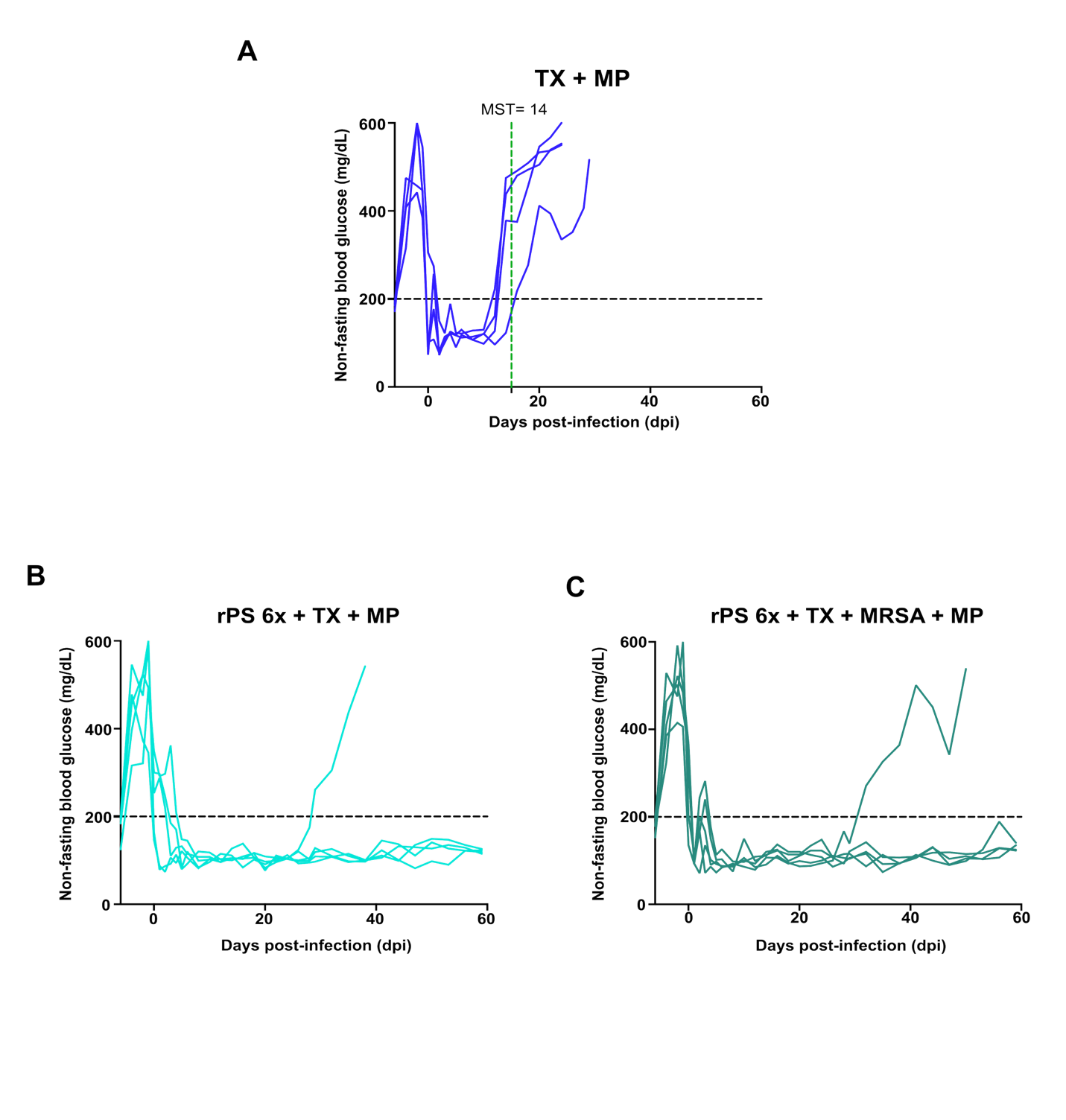

**Supplementary Figure S9: Blood glucose levels from -6 dpi to 59 dpi for treatment groups involving methylprednisolone (MP). (A)** MP + TX (N=4) with MST = 14 days. **(B)** rPS 6x + MP + TX (N=5). **(C)** rPS 6x + MP + TX + MRSA (N=5).

Supplemental materials & methods

*Materials*

All chemicals and reagents used in the polymer synthesis were purchased from Sigma-Aldrich unless stated otherwise.

*Polymer synthesis*

Poly(ethylene glycol)_17_-b-poly(propylene sulfide)_30_ (PEG_17_-*b*-PPS_30_) was synthesized as previously reported elsewhere.^9,19–21^ Briefly, methyl ether PEG (MW 750) was mesylated, converted to PEG-thioacetate, and activated with sodium methoxide to initiate ring-opening polymerization of propylene sulfide. Following polymerization, excess benzyl bromide was added as an end-capping agent to yield PEG_17_-PPS_30_. The degree of polymerization of the poly(propylene sulfide) (PPS) block was assessed via ^1^H NMR in CDCl_3_. Formation of the diblock copolymer was confirmed using gel permeation chromatography (GPC) with high performance liquid chromatography (HPLC) grade tetrahydrofuran as the mobile phase.

*Nanocarrier formulation and characterization*

rPS were formulated following the thin-film hydration method as previously described.^9,22^ Briefly, PEG_17_-PPS_30_ (20 mg) were dissolved in dichloromethane and rapamycin (1 mg; MedChemExpress) were dissolved in ethanol (25mg/mL), mixed together, dried as a thin film, and hydrated in PBS with shaking for 24h. Unencapsulated rapamycin was separated via size exclusion chromatography using Sephadex LH-20 followed by extrusion of the polymersomes through a 0.2 μm polyethersulfone (PES) membrane. Blank PS were prepared identically, omitting rapamycin addition. All formulations underwent extensive characterization via dynamic light scattering (DLS), cryogenic transmission electron microscopy (Cryo-TEM) and small-angle X-ray scattering (SAXS). DLS measurements were carried out on Zetasizer Ultra (Malvern) to assess the diameter of the nanocarriers and the polydispersity index of the formulation. Cryo-TEM (Thermo Fisher Glacios TEM 200 kV) and SAXS (X-ray beam with a wavelength of λ = 1.24 Å and an energy of 9 keV) were utilized to confirm the spherical architecture and presence of the bilayer organization. Rapamycin encapsulation was quantified by HPLC as previously described.^9^  Endotoxin was measured using the Chromogenic LAL Endotoxin Assay kit (GenScript) and only formulations with endotoxin levels < 0.05 EU/mL were considered for *in vitro* and *in vivo* testing.

*Neutrophil isolation, treatment and functional assays*

Murine bone marrow neutrophils were isolated by density-gradient centrifugation as previously described by Monteith et al.^23^ Cells were rested on ice for 45 min and equilibrated in D10 medium before 2h treatment with rapamycin, blank PS, or rPS (10 μg/mL), with polymer concentration matched between blank PS and rPS groups. Cells were stimulated with LPS (1 μg/mL) for ROS assays or live *S. aureus* JE2 (MOI 10) for NETosis and bacterial survival studies. For ROS production, neutrophils were incubated with H_2_-DCFDA (5 μM; ThermoFisher Scientific) and fluorescence was measured at 494/518 nm. For NETosis, neutrophils were cultured with *S. aureus* for 3 h, fixed, and stained for PECy7 anti-mouse CD11b (Tonbo, cat. 60-0112-U100), BV711 anti-mouse Ly6G (Biolegend, cat. 101219), myeloperoxidase (MPO) (Abcam, cat. ab90810), and citrullinated Histone H3 (H3Cit) (Abcam, cat. ab5103). Secondary staining was performed using Alexa Fluor 488 streptavidin (Biolegend, cat. 405235) and Alexa Fluor 647 anti-rabbit (Biolegend, cat. 406414). Suicidal NETosis was defined as dsDNA+MPO+H3Cit+ events among CD11b+ Ly6G+ neutrophils on flow cytometry.^24^ Parallel co-cultures were lysed for CFU enumeration to assess bacterial survival which was calculated as CFU recovered relative to the initial inoculum and expressed as percent survival. Neutrophil viability was determined by Ghost dye (Tonbo, cat. SKU 13-0863-T100) staining in unstimulated treatment groups.

*Mice*

Male C57BL/6, BALB/c, and C3H mice (8 weeks old; Jackson Laboratory) were housed at the Northwestern University Center for Comparative Medicine. Animals were housed in the biosafety level 2 facility and used in agreement with the Northwestern University Institutional Animal Care and Use Committee under protocols IS00029556 and IS00029483.

*Staphylococcus aureus culture*

For *in vitro* experiments*, Staphylococcus aureus* (strain JE2; a MRSA strain and USA300 isolate) was inoculated with 5 mL in tryptic soy broth and incubated overnight. The next day, the bacteria was opsonized at 1 to 10 dilution in a 1:1 mixture of DMEM + FBS for a minimum of 30 minutes before use in the experiments. For *in vivo* experiments, MRSA was cultured in tryptic soy broth supplemented with erythromycin (5 μg/mL) at 37°C under agitation for 5h to yield MRSA in the logarithmic phase. Optical density measurements (OD_600_) were used to determine viable bacterial titers. Bacteria were pelleted and resuspended at the desired concentration in PBS for *in vivo* experiments.

*Bacterial burden, cytokine and MPO quantification*

Organ homogenates were serially diluted and plated on tryptic soy agar containing erythromycin for CFU enumeration, with values normalized to tissue mass. Skin samples were sonicated in an ice bath before plating. Serum and skin homogenates (100 mg/mL tissue) cytokines were measured using the BD Cytometric Bead Array Mouse Th1/Th2/Th17 kit (BD Biosciences) according to the manufacturer’s instructions. Collected data were analyzed using the FCAP Array Infinite Software (SoftFlow). MPO activity was quantified in serum and skin homogenates using the Myeloperoxidase Activity Assay kit (Invitrogen, cat. EEA016), following the manufacturer’s instructions.

*Flow cytometry of the spleen in immunotherapy + MRSA infection study*

Splenocytes were prepared by mechanical dissociation through 70-μm strainers followed by red blood cell lysis. Cells were stained with Zombie NIR Biolegend, cat.77184), blocked with anti-mouse CD16/CD32 (TruStain FcX; BioLegend, cat. 101319), and stained with the following surface antibodies: BV750 anti-mouse CD45 (Biolegend, cat.103157), BV785 anti-mouse/human CD11b (Biolegend, cat.101243), Pacific Blue anti-mouse CD11c (Biolegend, cat. 117322), Alexa Fluor 700 anti-mouse Ly-6G (Biolegend, cat. 127622), PerCP-eFluor 710 anti-mouse CD64 (Invitrogen, cat. 46-0641-82), BV510 anti-mouse CD3 (Biolegend, cat. 100234), PE/Cy5 anti-mouse CD4 (Biolegend, cat. 100410), BV421 anti-mouse CD86 (BD Horizon, cat. 564198), BV650 anti-mouse CD8a (Biolegend, cat. 100742), and APC anti-mouse CD19 (Biolegend, cat. 115512). Cells were then fixed and permeabilized using the Foxp3 Transcription Factor Staining Buffer Set (Invitrogen, cat. 00-5523-00) and stained intracellularly with PE anti-mouse IFN-γ (Biolegend, cat. 505808), PE/Cy7 anti-mouse IL-4 (Biolegend, cat. 504117), FITC anti-mouse IL-17A (Biolegend, cat. 506908), and eFluor 660 anti-mouse Foxp3 (Invitrogen, cat. 50-5773-82). Samples were acquired on a 3-laser Aurora cytometer (CyTek), unmixed in SpectroFlo, and analyzed in FlowJo v10 using FMO controls for gating. The gating strategy is described in Supplementary Fig. S3.

*Complete blood count (CBC) analysis*

Whole blood obtained via retro-orbital bleeding were collected in EDTA tubes and analyzed on an Element HT5 hematology analyzer providing a comprehensive panel, including white blood cells (WBC), lymphocytes, monocytes and neutrophils counts. The instrument also provides reference ranges for all given parameters.

*Methylprednisolone (MP) combination therapy study*

Experiments involving methylprednisolone (purchased from Northwestern University Pharmacy) followed the same transplantation and MRSA infection protocol described above, except that some recipients received a single intraperitoneal injection of MP (20 mg/kg) on the day of MRSA infection. Additional treatment groups included MP alone and MP combined with rPS.

*Statistical analysis*

Statistical analysis was conducted using GraphPad Prism version 10.5.0. Statistical tests were performed using unpaired 2-tailed t-tests, a one-way or two-way ANOVA with Tukey's multiple comparison post-test, and a Log-rank test (Mantel Cox) with Holm-Šídák post-hoc analysis for multiple survival curves comparison. Statistical significance was set at *P* < 0.05.
